# Fine-scale population structure and demographic history of British Pakistanis

**DOI:** 10.1101/2020.09.02.279190

**Authors:** Elena Arciero, Sufyan A. Dogra, Massimo Mezzavilla, Theofanis Tsismentzoglou, Qin Qin Huang, Karen A. Hunt, Dan Mason, David A. van Heel, Eamonn Sheridan, John Wright, Neil Small, Shai Carmi, Mark M. Iles, Hilary C. Martin

**Affiliations:** Wellcome Sanger Institute; University of Trieste; Bradford Institute for Health Research, Bradford Teaching Hospitals National Health Service (NHS) Foundation Trust, Bradford; University of Leeds; Blizard Institute, Barts and The London School of Medicine and Dentistry, Queen Mary University of London, London; Queen Mary University of London; Bradford Institute for Health Research, Bradford Teaching Hospitals National Health Service (NHS) Foundation Trust, Bradford BD9 6RJ, UK; Braun School of Public Health and Community Medicine, The Hebrew University of Jerusalem, Jerusalem, Israel; Wellcome Trust Sanger Institute

## Abstract

Previous genetic and public health research in the Pakistani population has focused on the role of consanguinity in increasing recessive disease risk, but little is known about its recent population history or the effects of endogamy. Here, we investigate fine-scale population structure, history and consanguinity patterns using genetic and questionnaire data from >4,000 British Pakistani individuals, mostly with roots in Azad Kashmir and Punjab. We reveal strong recent population structure driven by the *biraderi* social stratification system. We find that all subgroups have had low effective population sizes (N_e_) over the last 50 generations, with some showing a decrease in N_e_ 15-20 generations ago that has resulted in extensive identity-by-descent sharing and increased homozygosity. Using new theory, we show that the footprint of regions of homozygosity in the two largest subgroups is about twice that expected naively based on the self-reported consanguinity rates and the inferred historical N_e_ trajectory. These results demonstrate the impact of the cultural practices of endogamy and consanguinity on population structure and genomic diversity in British Pakistanis, and have important implications for medical genetic studies.

## Introduction

Estimates suggest that around 10% of the world’s population are offspring of closely related parents, mostly in north and sub-Saharan Africa, the Middle East, and west, central, and south Asia ^1^. In these regions, endogamy frequently co-occurs ^2^. Exploring the impact of consanguinity, endogamy, and population structure on genetic variation within these populations is important for quantifying their relative effects on risks of genetic diseases. Here, we investigate fine-scale population structure, history and consanguinity patterns in the British Pakistani population, one of the largest ethnic minorities in the United Kingdom, with a population size of 1.17 million in the 2011 census ^3^. The most substantial wave of immigration from Pakistan to the UK occurred in the 1950s/60s, after the partition of British India, with many Pakistanis coming to work in the steel, textile, and engineering industries, or as doctors, followed later by family members ^4,5^. British Pakistanis (i.e. individuals with Pakistani ancestry born either in Pakistan or in the UK) are one of the most socioeconomically disadvantaged ethnic groups in Britain ^6^. They have rates of type 2 diabetes and heart disease that are 2-4 times higher than the White British population ^7–9^, as well as an increased risk of congenital anomalies due to the prevalence of consanguinity ^10^. These factors, combined with the laudable drive to increase the number of genetic studies on people with non-European ancestry ^11^, have spurred the creation of several cohort studies whose aims include exploring the environmental and genetic contributions to various phenotypes in Pakistani-ancestry individuals and the impact of homozygous gene knockouts ^12^. These include Genes & Health (G&H) ^13^ and Born in Bradford (BiB) ^14^ (in the UK), and the Pakistan Risk of Myocardial Infarction Study (PROMIS) ^15^. A proper understanding of the demographic history of the target populations is important for designing robust and effective genetic analyses, as has been recently highlighted in work on polygenic scores ^16,17^.

Modern South Asians are a mixture of different proportions of two ancestral populations: the Ancestral Northern Indian (ANI) and Ancestral Southern Indian (ASI) components ^18,19^. Northwest Indians and Pakistanis have a greater proportion of the ANI component ^18,19^. Several major studies of human genetic diversity, such as the Human Genome Diversity Project (HGDP) ^20^, 1000 Genomes ^21^ and GenomeAsia ^22^, have highlighted differences between multiple ethnic groups within Pakistan, as well as the elevated levels of autozygosity due to consanguinity ^23^. These previous studies have focused mostly on population structure on a macro-scale, with relatively limited sample sizes per population, and they lacked information on finer-scale groupings within each of these populations.

Most of the ethnic and tribal groups in Pakistan follow a patrilineal kinship system based on the *biraderi* (brotherhood) that shares its historical roots with the better-studied Indian caste system. The *biraderi* system is a means of attributing social status and providing mutual social support ^24^. Some *biraderi* groups such as Rajput and Jatt have been present on the Indian sub-continent for the last 2,000-3,000 years ^25–27^. Other *biraderi* originated in the early Medieval period, such as the Gujjars whose identity is traced back to the Gurjara kingdom in present-day Rajasthan around 570 CE ^28^. Mass-conversion to Islam during the pre-Mughal and Mughal era introduced multiple new *biraderi*, including Syed, Qureshi, Malik and Sheikh ^29^. Historical records suggest that endogamous practices were strengthened during the Mughal Empire to ensure order in society ^30^, and became even stricter during the colonial times of the 19th century, as the social classification system was reinforced by the British to solidify their political authority, enable rationalised taxation, and establish rules about individual and family property ^30,31^. Overall, however, there are limited historical records about when the *biraderi* groups emerged or the extent to which endogamy was practiced over the centuries. Very little is known about the effect of this historical social structure on present-day genetics, and previous work has been limited to small studies of a few microsatellite markers ^27,32,33^.

Here, we analyse genotype array data and exome sequence data from thousands of Pakistani-ancestry individuals sampled in Britain as part of the Born in Bradford (BiB) project. BiB is a birth cohort set up to investigate the social, environmental, and genetic causes of poor health and educational outcomes of children born in Bradford, a city with high levels of socio-economic deprivation in the north of England ^14^. Around half of the individuals in this cohort have Pakistani ancestry, coming primarily from Azad Kashmir (Mirpur) and northern Punjab (see map in Supplementary Figure 1), in proportions similar to the British Pakistani community as a whole ^5,35^. BiB has rich self-reported information on the Pakistani mothers’ *biraderi* groups, places of birth, and parental relatedness. To our knowledge, this is the largest sample of Pakistani-ancestry individuals analysed to date to explore population genetic questions.

## Results

### Samples and dataset

We assembled a large dataset of 7,180 individuals with Pakistani ancestry (<u>Supplementary Tables 1 and 2</u>) from the BiB project, of which 5,669 had been genotyped on the Illumina CoreExome array and 1,511 on the Illumina Global Screening Array (GSA); 2,484 of these also had exome-sequence data. Identifying related individuals is challenging in a population with high consanguinity and endogamy, so we tried several algorithms (Methods, Supplementary Figure 2). We erred on the side of caution and used the algorithm that gave the highest estimates of kinship, PropIBD from KING ^36^, to identify and remove putative relatives (3rd degree or closer). Most analyses in this paper are based on 2,200 unrelated mothers genotyped on the CoreExome array, with 1,616 unrelated children (CoreExome) and 228 unrelated fathers (GSA) used in some analyses.

After cleaning the questionnaire responses about *biraderi* membership, we determined that fifty-six distinct groups had been reported. Most of these are recognised *biraderi* groups, but some individuals identified themselves with tribal/regional groups (e.g. Pathan, or Kashmiri), or clans within a *biraderi* (e.g. Choudhry) (<u>Supplementary Table 3</u>). Presuming that individuals have reported the labels that best represent their group identity within the context of the Bradford Pakistani community, we henceforth refer to these collectively as ‘subgroups’ rather than ‘*biraderi*’.

### Genetic diversity of Bradford Pakistanis in a worldwide context

We first investigated the genetic relationships between the Bradford Pakistanis and other worldwide populations from publicly available datasets (HGDP, Human Origins and 1000 Genomes). Principal component analysis (PCA) ^37,38^ showed that Pakistanis from Bradford cluster with other South Asians (Supplementary Figure 3a). Focusing on Pakistani groups, the Bradford Pakistanis lie between the Sindhi, Pathan and Burusho (Supplementary Figure 3b,c), as expected given they are predominantly from Punjab and Kashmir (Supplementary Figure 1). Identification of ancestral components using ADMIXTURE ^38^ (Supplementary Figure 4a) showed that Bradford Pakistanis have a similar genetic profile to other South Asian populations, and display little variation in their ancestral components in this broad context. Only the Pathans stand out; they have a higher fraction of the pink and blue components seen in Europeans. This is consistent with the fact that they represent a distinct ethnic group from Punjabis and Kashmiris, speaking a language from a different family (Pashto from the Iranian family, as opposed to Punjabi from the Indo-Aryan family). Outgroup *f3* statistics^39^ (Supplementary Figure 4b) confirmed genetic affinity between the BiB Pakistanis and the HGDP Pathans, 1000 Genomes Punjabis, and northern (Uttar Pradesh Brahmins) and western (Kashimiri Pandit) Indian populations. They also have genetic similarity to Central Asia populations (Supplementary Figure 4a,b), consistent with previous studies that have demonstrated the ANI component in modern Pakistanis ^18,19^.

We next investigated whether the self-reported subgroups who claim to have recent Arabic ancestry had a different pattern of genetic sharing with Middle Eastern populations compared to the other subgroups. We did not find statistically significant differences between groups using outgroup *f3* or *f4* statistics (Supplementary Figure 4c, Supplementary Table 4). We also saw no differences in the distribution of Y chromosome haplogroups between the *biraderi* that report Arabic ancestry and the other subgroups. Eighty-nine percent of individuals belong to the IJ* Y haplogroups, which are prevalent in Pakistan and Central Asia ^40,41^, while the rest have other haplogroups that are present in Central and South Asia (<u>Supplementary Table 5</u>)^42,43^. Similarly, the mothers have mitochondrial haplogroups that are common in Central and South Asia ^44,45,46,47,48^ (<u>Supplementary Table 6</u>).

### Population structure

We next investigated the fine-scale population structure within the Bradford Pakistanis. PCA of the samples revealed clear structure, with the first three principal components (PCs) each explaining ~4% of the variation and driven respectively by the separation of the Jatt and Choudhry subgroups, the Pathan, and the Bains and a subset of the Rajput individuals (Figure 1). The fact that the Choudhry and Jatt subgroups cluster together is consistent with the fact that “Choudhry” is an honorary title in Punjab and Kashmir used most commonly by the Jatts, who are the one of the largest ethnic groups in Pakistan and north-west India ^27^. ADMIXTURE analysis of the Bradford Pakistani samples alone indicates that the subgroups that were distinguishable on the PCA have different proportions of genetic components (Figure 1c). The Rajput group contains two distinct subgroups that we term henceforth Rajput-A and Rajput-B; the Rajput-B group, which has a higher fraction (> 40%) of the “red” ancestry component, appears more similar to Bains than to Rajput-A, and these individuals cluster with Bains on the PCA (Figure 1b). We found no evidence that the Rajput-A versus Rajput-B distinction is driven by geographical origin within Pakistan (Supplementary Figure 5). It may well be that Rajput-B and the different subgroups of Rajput-A represent different sub-clans of Rajputs, of which there are hundreds across South Asia ^49,50^. An alternative explanation is that some people self-identifying as Rajputs actually have diverse ancestries since individuals from other subgroups chose to identify with this group to benefit from land allocations during British colonial times ^24,51^ or to increase their social status after migration to the UK ^52^. The fact that Bains and the Rajput-B group appear to form a homogeneous genetic cluster is consistent with historical evidence that Bains is one of the Rajput families ^53^.

**Figure 1:**
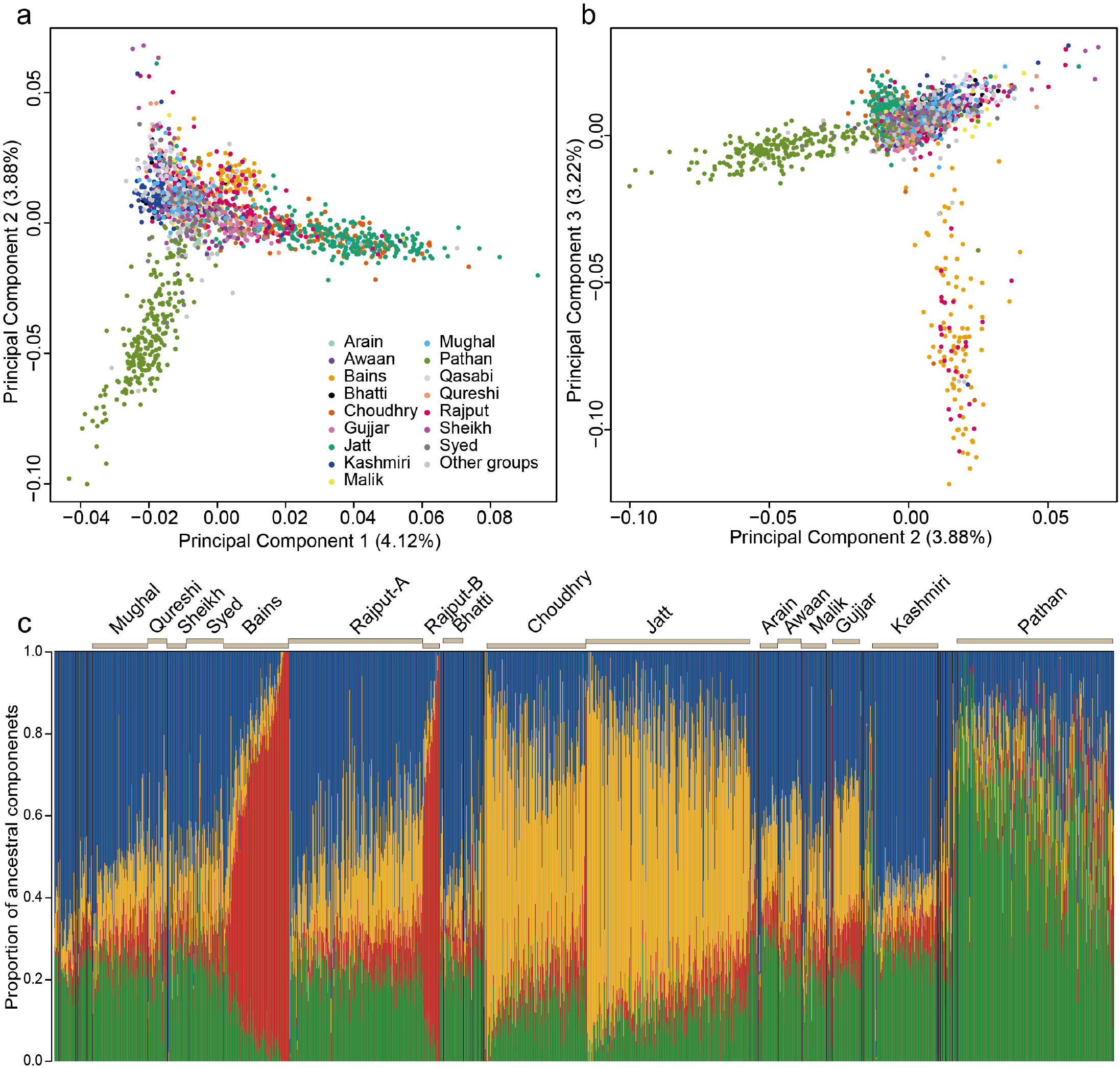
Population structure within British Pakistanis from Bradford. (a,b) Principal components analysis of 2,200 unrelated Pakistani mothers, with the self-reported subgroups with >20 samples indicated in different colours. Plots show PC1 versus PC2 (a) and PC2 versus PC3 (b). Proportion of overall variation explained by each PC is noted in brackets on the axis label. (c) ADMIXTURE plot (K=4) illustrating different ancestral components making up the various subgroups, with the largest subgroups indicated. We have indicated the two distinct subgroups amongst the Rajput.

We next applied Uniform Manifold Approximation and Projection (UMAP) to the first 20 PCs (Supplementary Figure 6). This allowed us to define the Pathan and the Bains/Rajput-B subgroups more cleanly than on the PCA, but failed to distinguish additional groups. We found no significant correlation (Mantel test p-value: 0.9384) between genetic distance (measured by UMAP1 and UMAP2 vectors) and geographic distance (measured by geographic coordinates of the individual’s or her parents’ self-reported village of origin in Pakistan).

We then applied fineSTRUCTURE ^54^, a Bayesian clustering algorithm, to a matrix of haplotype-sharing patterns. The inferred hierarchical clustering tree based on 1,520 individuals from the sixteen major subgroups (Figure 2, Supplementary Figure 7) identified three clusters containing the majority of the self-reported Pathan (Cluster 8), Bains and Rajput-B (Cluster 9), and Jatt and Choudhry (Cluster 10) individuals. Subsets of Bains and Rajput-B (Cluster 6) and Jatt and Choudhry (Cluster 11) clustered with individuals from other groups, although Cluster 11 had lower support than the other clusters (Figure 2). The Bains and Rajput-B in Cluster 6 showed a smaller proportion of the red component in the ADMIXTURE plot (Figure 1c) than those in Cluster 9 (Wilcoxon rank sum test p-value=1×10^−12^). For other self-reported groups, the majority of individuals from the group fell in a single fineSTRUCTURE cluster: the Kashimiri (Cluster 1), Syed and Awaan (Cluster 2), Arain (Cluster 4), Gujjar (Cluster 5), and Qasabi (Cluster 7). Some of the self-reported groups appear to be quite heterogeneous, with individuals from these groups distributed across different inferred clusters; Rajput-A is a notable example (Figure 2). Rerunning fineSTRUCTURE a second time and running it including the minor subgroups (Supplementary Figure 9) produced a very similar tree. Genetic differentiation was very low (F_ST_< 0.001) between the Awaan and Syed in Cluster 2, the Bains and Rajput-B in Cluster 9, and the Jatt and Choudhry in Cluster 10 (Supplementary Figure 8, <u>Supplementary Table 7</u>). We henceforth pooled together individuals in these very similar subgroups who fell within those dominant clusters (Awaan/Syed, Bains/Rajput-B, and Jatt/Choudhry), and restricted the Arain, Gujjar, Kashmiri, Pathan and Qasabi subgroups to those individuals who fell within the dominant fineSTRUCTURE cluster for that group.

**Figure 2:**
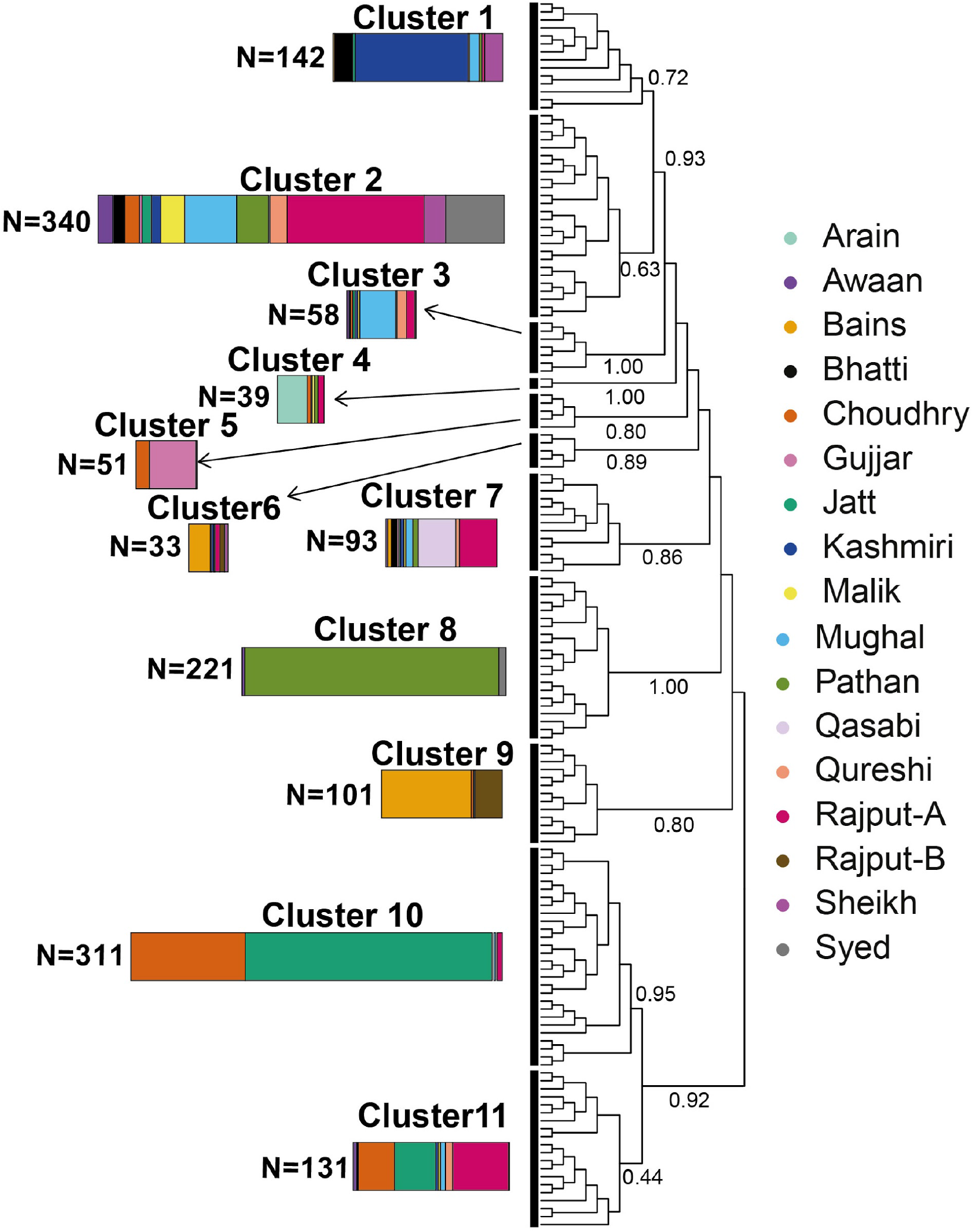
Fine-scale population structure inferred amongst 1,520 Bradford Pakistani mothers using fineSTRUCTURE. The tree illustrates the results of hierarchical clustering of the co-ancestry matrix using patterns of haplotype sharing from ChromoPainter. Each ‘leaf’ on the tree contains multiple individuals. The coloured bars represent the composition of each of the major clusters indicated by the thick black vertical lines. The length of each coloured bar is proportional to the number of individuals in that cluster, with the proportion of each colour representing the fraction of individuals from each self-reported subgroup that make up the cluster. The labels on the edges are the posterior assignment probabilities from fineSTRUCTURE. Only the individuals from the sixteen largest groups were included here, but several other smaller subgroups could be distinguished when applying fineSTRUCTURE to all 2,220 unrelated mothers (Supplementary Figure 9).

The *biraderi* system is culturally characterised by patrilineal kinship ties, so we examined GSA data on 228 unrelated Pakistani fathers from BiB to test whether males from the same subgroups did indeed carry the same Y haplogroups (<u>Supplementary Table 5</u>). Despite the clear delineation of some groups observed through analysis of the autosomal data, we found that even males within the most distinct groups (e.g. Jatt/Choudhry, Bains/Rajput-B, Pathan) did not tend to cluster together by Y haplogroup (Supplementary Figure 10). This is consistent with previous findings in Punjabi Rajputs ^33^ and in Jatts ^27^, and may suggest that the founders of each *biraderi* included males with several different haplogroups, or that, historically, the patrilineality of the *biraderi* system was not very strict.

### Demographic history, endogamy and consanguinity

We next estimated population divergence times between the homogeneous subgroups defined in the previous section, using the NeON package ^55^ which considers patterns of linkage disequilibrium decay. We found that all groups diverged from one another within the last ~70 generations (Figure 3a, <u>Supplementary Table 8</u>, <u>Supplementary Table 9</u>), consistent with the low genetic differentiation between the subgroups (Supplementary Figure 8). The estimates in Figure 3a suggest that the history of these subgroups cannot be considered as a series of clean splits between ancestral populations; rather, it appears that several groups began to differentiate from one another around the same time, with some degree of admixture persisting between the groups after their initial formation. Nonetheless, we can see that Bains/Rajput-B (Cluster 9 in Figure 2) show the oldest divergence time from the other groups, followed by Qasabi (Cluster 7) and Pathan (Cluster 8) (<u>Supplementary Table 8</u>). Within clusters, Bains and Rajput-B (Cluster 9) have a divergence time estimate close to zero, as do Jatt and Choudhry in (Cluster 10; <u>Supplementary Table 9</u>), consistent with their clustering together on fineSTRUCTURE and having very low F_ST_ (F_ST_ < 0.001).

**Figure 3:**
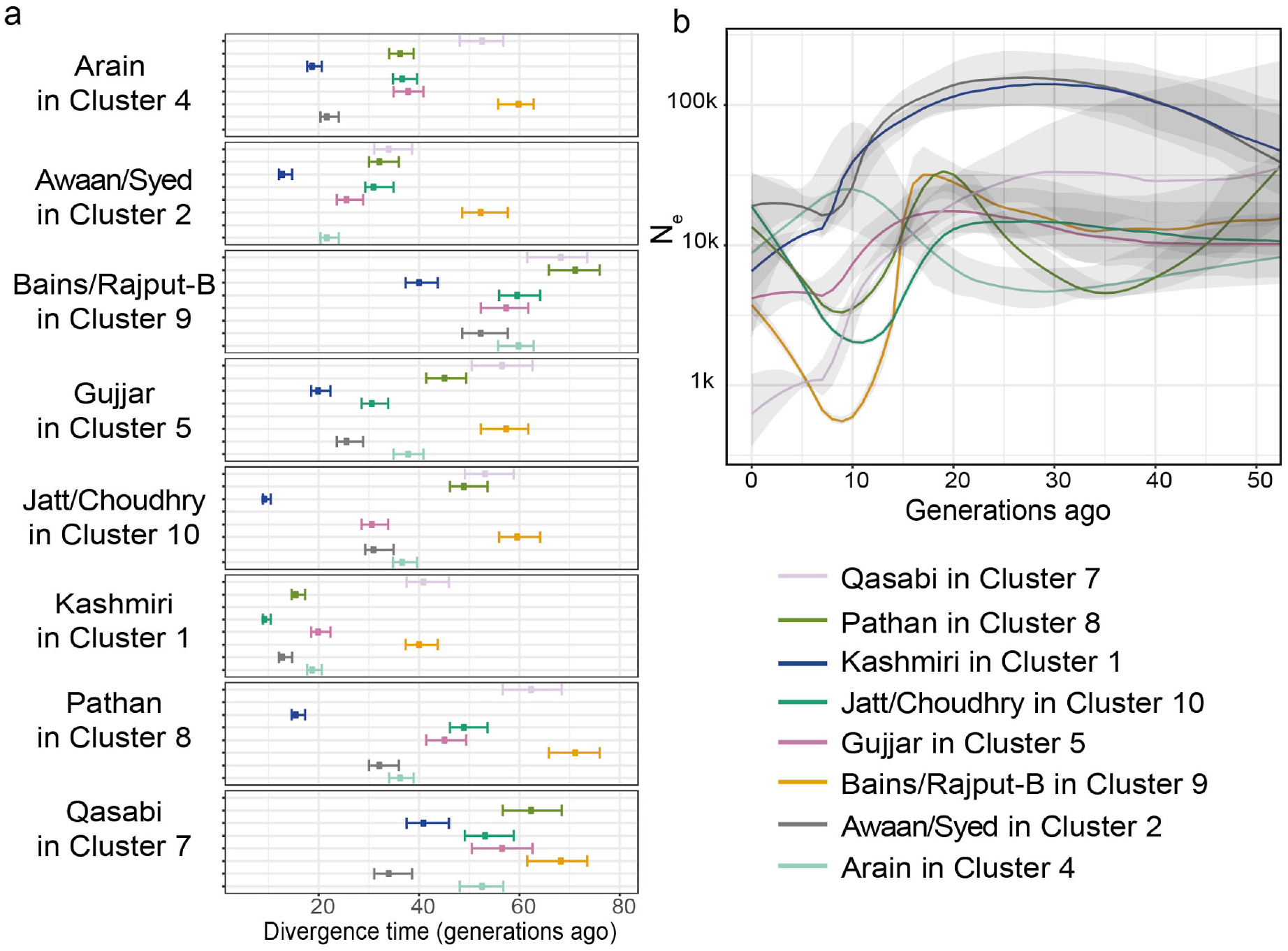
Divergence times and historical effective population size changes of Bradford Pakistani subgroups. Note that these are the homogeneous subgroups defined using the fineSTRUCTURE clusters. (a) Divergence times estimated using NeON ^55^. Within each panel, the points show the estimated divergence time between the group indicated on the left on the y-axis and those indicated in the legend. The horizontal lines indicate 95% confidence intervals. (b) Change in effective population size (N_e_) through time estimated with IBDNe. The coloured lines indicate the mean estimate and the grey shading indicates 95% confidence intervals. Supplementary Figure 13b shows results from IBDNe using the fineSTRUCTURE clusters.

Population relationships and gene flow inferred by Treemix ^56^ suggested that the Bains/Rajput-B group is characterized by the strongest genetic drift, followed by Qasabi and Pathan (Supplementary Figure 11). The tree topology without migration edges explained most of the variance (97.4%), but adding one or two migration edges increased the variance explained to >99%, detecting gene flow events from Pathan into Kashimiri (one migration edge), and from Bains/RajputB and Jatt/Choudhry into Kashmiri (two migration edges) (p-values<0.001). *f3*-statistics ^39^ also showed that the Kashmiri were characterised by significant admixture events with other subgroups (Supplementary Figure 12, <u>Supplementary Table 11</u>). Although about half of the Bradford Pakistanis come from Azad Kashmir, people who refer to themselves as “Kashmiri” are believed to originate specifically from the Kashmir Valley ^57^. Our finding that the Kashmiri have gene flow from other groups is consistent with historical evidence that the Kashmiri from the valley belonged to different *biraderi* and tribes. Some migrated to Azad Kashmir and Punjab during the 19th century, where they began identifying themselves as Kashmiri ahead of their *biraderi* identity, tending to marry within this group ^53,58,59^. This likely explains the relative genetic homogeneity of the Kashmiri in our sample (80% of individuals fall in Cluster 1 on Figure 2) and the observation that the most recent estimated divergence time between Kashmiri and other groups is ~10 generations ago (Figure 3a).

We next applied IBDNe ^60^ to infer recent effective population sizes (N_e_) through time across the different subgroups. All subgroups were inferred to have relatively low N_e_ over the last 50 generations compared to white British individuals (Supplementary Figure 13a), likely reflecting the endogamy of the *biraderi* system. Bains/Rajput-B (Cluster 9), Jatt/Choudhry (Cluster 10) and Pathan (Cluster 8) showed a similar trend: a strong reduction in N_e_ around 10-20 generations ago followed by a recovery (Figure 3b). The other subgroups and clades, except for Arain, showed a progressive decrease in N_e_ in the last 15 generations (Figure 3b), which may reflect the increase in endogamy that occurred during the Mughal Empire and British colonial times ^30^.

To quantify the extent of this historical endogamy, we calculated identity-by-descent (IBD) scores ^34^ in the Pakistani subgroups. These scores represent the average total length of IBD shared between any two individuals from the same subgroup after excluding relatives, considering segments between 5 and 30 cM (Supplementary Figure 14). Most of the Pakistani subgroups have substantially higher IBD scores than other worldwide populations from the 1000 Genomes Project, including the Finns (Figure 4a), similar to those reported previously for some isolated Indian groups ^34^. The Bains/Rajput-B group had the highest IBD score, followed by Qasabi and Jatt/Choudhry. Similar results were obtained when constructing IBD scores using the same length filtering as Nakatsuka *et al*. ^34^, and when focusing on the fineSTRUCTURE clusters (Supplementary Figure 15).

**Figure 4:**
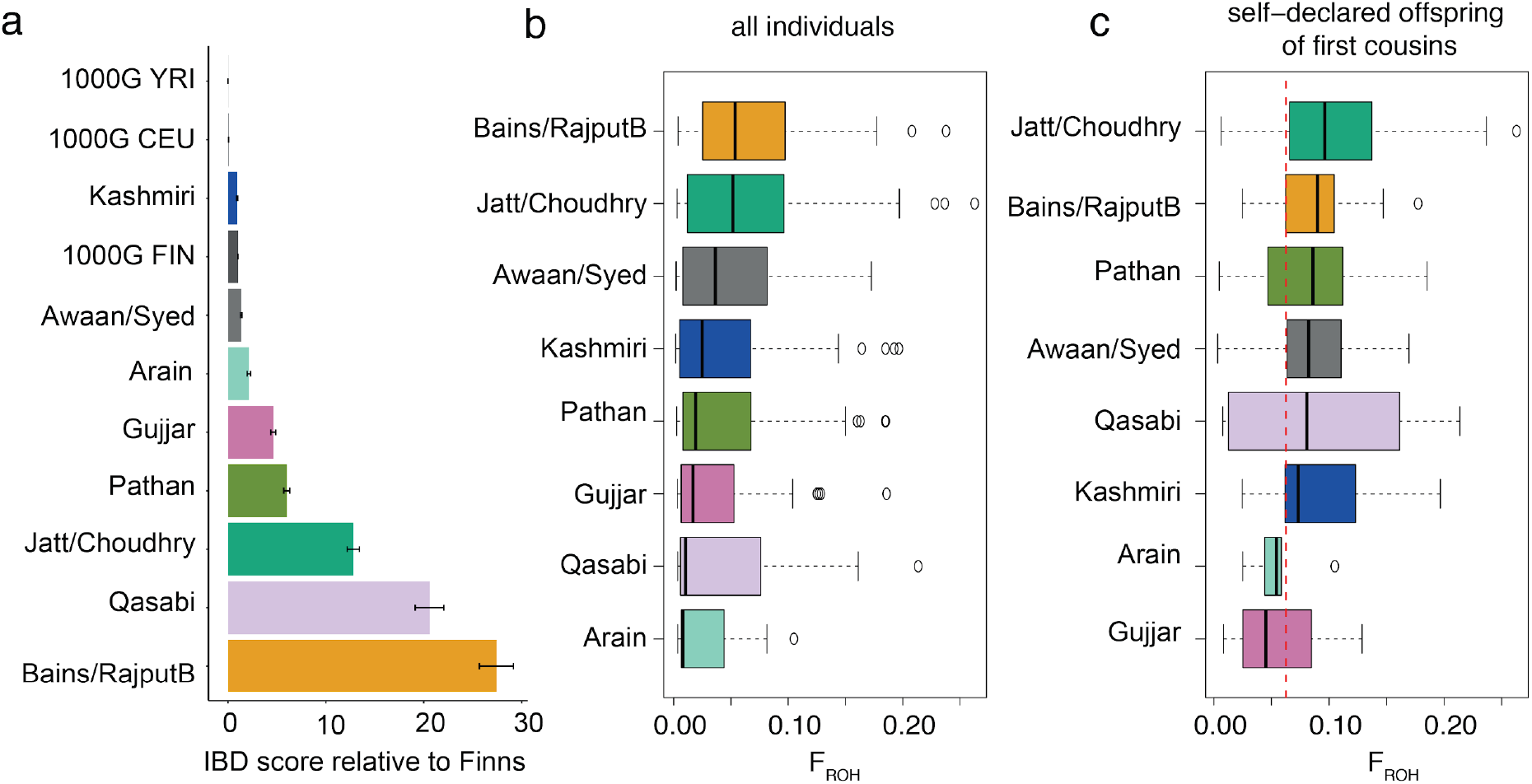
Effect of endogamy and consanguinity on IBD sharing and ROH patterns in BiB Pakistani subgroups. Note that these are the homogeneous subgroups defined using the fineSTRUCTURE clusters. a) IBD scores calculated as the average total length of IBD segments between 5 and 30cM shared between individuals from the indicated group, standardised by the value for the 1000 Genomes Finns (FIN). The error bars indicate standard errors. YRI: Yoruba; CEU: Western Europeans. b) Boxplots showing the distribution of the fraction of the genome in regions of homozygosity (F_ROH_) broken down by subgroup. (See Supplementary Figure 16 for a breakdown by self-reported group.) c) As for b) but with only the individuals who self-reported as being offspring of first cousins. The red vertical line indicates the expectation (1/16) for offspring of first-cousins whose parents are themselves unrelated.

Fifty-seven percent of the BiB Pakistani mothers reported that their parents were related, and 63% reported being related to their child’s father (<u>Supplementary Tables 12</u> and <u>13</u>). As expected, a much higher fraction of the genome was homozygous (F_ROH_) in the Pakistani mothers than the White British (Supplementary Figure 16). Amongst the BiB children, those whose parents were both born in the UK had significantly lower F_ROH_ than those whose parents were both born in Pakistan, and both groups had significantly lower F_ROH_ than those who had one parent born in Pakistan and the other in the UK (Supplementary Figure 17a). This likely reflects changing marriage preferences in second-generation British Pakistanis ^61^. F_ROH_ differed significantly between the subgroups (ANOVA; F-test p=8×10^−8^; Figure 4b). In a joint model, self-reported consanguinity explained 41% of the variance in F_ROH_ (ANOVA F-test p<1×10^−173^), self-reported subgroup explained 2% (p=0.017) and their interaction explained 6% (p=0.016). Hence, while both consanguineous marriage and *intra*-biraderi marriage make a significant contribution to homozygosity, the former explains more variance. The pattern of differences in F_ROH_ across subgroups was similar between individuals who were born in Pakistan and those born in the UK (Supplementary Figure 17b), implying that each subgroup’s *relative* frequency of consanguineous marriage has not changed substantially after migration. We saw variation between subgroups even amongst individuals who reported that their parents were first cousins (ANOVA; F-test p=0.01; Figure 4c). This likely partially reflects levels of consanguinity in previous generations, and it is notable that the groups with the highest homozygosity amongst offspring of first cousins (Jatt/Choudhry, Bains/Rajput-B) also have higher levels of IBD sharing (Figure 4a). For many of the subgroups, long regions of homozygosity (ROHs) between 5 and 50 cM are seen even in individuals who report that their parents are unrelated (Supplementary Figure 18), particularly in the Bains/Rajput-B group. It may be that there is under-reporting of close consanguinity, but low N_e_ due to endogamy is also undoubtedly contributing to the landscape of ROHs as well as to IBD sharing between individuals.

To model the joint effects of consanguinity and endogamy on ROHs and IBD segments, we applied recently developed theory ^62^ to predict their length distribution given historical consanguinity rates and historical N_e_ trajectory. We focus on the two largest homogeneous subgroups: the Pathan from Cluster 8 (N=213) and the Jatt/Choudhry from Cluster 10 (N=299) (Figure 2). As an initial naive estimate of consanguinity rate, we used the proportion of mothers in each group who reported that their parents were first cousins, second cousins, or ‘other blood/other relative’, which we assumed were third cousins. This gave an estimate of the average kinship between spouses of 0.016 for Pathan and 0.028 for Jatt/Choudhry (see Methods). Supplementary Figure 19 illustrates the importance of considering not only average kinship between spouses but also historical N_e_ when evaluating the proportion of the genome covered by ROHs of different lengths (the ‘ROH footprint’); the N_e_ has a major influence on the expected footprint of ROHs < 25cM, but much less on ROHs longer than this, since those are primarily due to mating between close relatives. We ran the model from ^62^ using the N_e_ estimates from IBDNe (Figure 3b) as input, varying the average spousal kinship (see Methods). Figure 5 compares the observed ROH and IBD footprints to those expected under the models with different parameters. It indicates that the observed ROH footprint determined with bcftools/roh is not consistent with the average spousal kinship we estimated naively from the parental relationships reported by the mothers, but is about twice as high as this: ~0.035 in the Pathans and ~0.056 in the Jatt/Choudhry group. This likely reflects the influence of consanguinity in earlier generations on the realised relatedness between spouses (Supplementary Figure 2c,d). Similar results were obtained with ROHs called by PLINK, whereas GARLIC seems to call more short ROHs than the model predicts (Supplementary Figure 20), consistent with the higher false positive rate for short ROHs reported in the original GARLIC paper ^63^.

**Figure 5.**
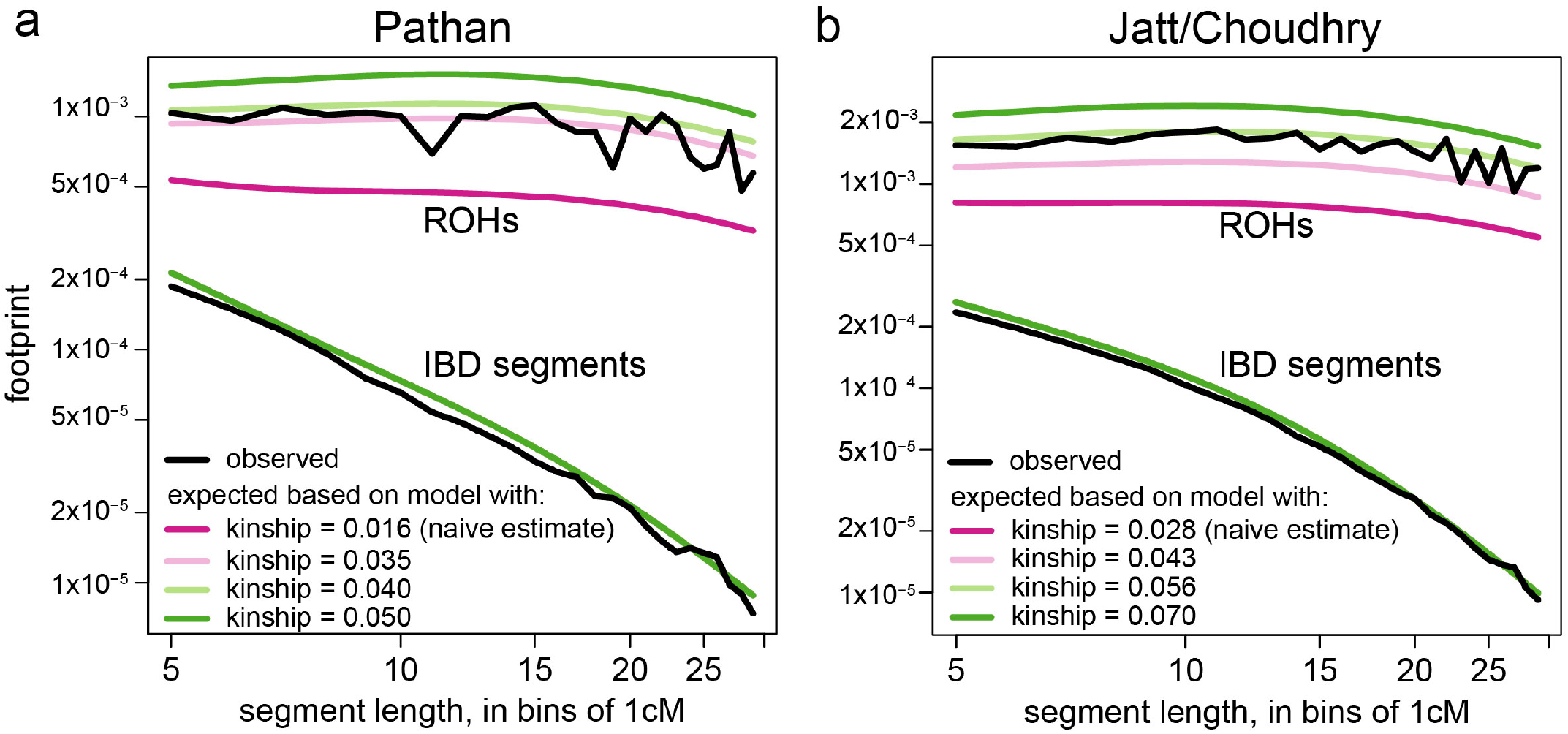
Observed ROH and IBD footprints compared to the expectation under a coalescent model^62^. The footprint is the average fraction of the genome covered by segments of a given length interval. The top lines represent the ROH footprint and the bottom lines the IBD footprint. Points are plotted at the beginning of each 1cM interval. The N_e_ profile used to compute the expected footprints was determined by IBDNe using the Pathan from fineSTRUCTURE’s Cluster 8 (a) or Jatt/Choudhry from Cluster 10 (b). We used several different values for the average kinship between spouses (see Methods), as indicated by the green and pink lines. Note that for IBD segments, the expectation is unaffected by the kinship value. The observed footprints (black lines) were determined using filtered IBD and ROH calls from IBDseq and bcftools/roh respectively. Supplementary Figure 20 shows equivalent plots using the upper and lower bounds of the 95% confidence interval for the IBDNe estimate as the N_e_ trajectory, and using observed footprints from different ROH or IBD calling/filtering strategies.

### Searching for founder effects

A reduction in N_e_ due to endogamy can lead to founder effects, which may push rare pathogenic variants to higher frequency ^34,64^. To investigate this, we searched for variants with significant allele frequency differences between each of the three largest homogeneous Pakistani subgroups (Pathan in Cluster 8, Bains/Rajput-B in Cluster 9, Jatt/Choudhry in Cluster 10) and all individuals in Clusters 1 to 5 and Cluster 7 (Figure 2). We used three approaches: pcadapt ^65^ and GEMMA-LMM ^66^ for outlier detection on the CoreExome genotype data, and a simple Fisher’s exact test on variants ascertained with whole-exome sequencing data, following previous work ^67^ (see Methods). We found 215 significant variants in 140 independent loci (r^2^<0.2) after multiple testing correction on at least one of these tests (Supplementary Figure 21, <u>Supplementary Tables 14</u> and <u>15</u>), including a total of forty-eight protein-altering variants. A few of these were in Mendelian disease-causing genes, but we found no evidence that these were likely to be pathogenic.

We noted multiple variants in the HLA region that had significantly different frequencies in the Jatt/Choudhry subgroup (Supplementary Figure 21), several of which are significantly associated with various complex traits such as leprosy (rs9274741; p=4×10^−10^ in ^68^), IgA nephropathy (rs9275596; p=3×10^−31^ in ^69^), idiopathic membranous nephropathy (rs3115663, rs3130618, rs3134945, rs11229; p<2×10^−40^ in ^70^), complement C4 levels (rs2071278; p=4×10^−72^ in ^71^) and haemoglobin levels (rs10885; p=2×10^−36^ in ^72^). Variants in the HLA are known to have been subject to selection in multiple populations, likely because they confer resistance to pathogens ^73^. It might be that this signal in the HLA in the Jatt/Choudhry subgroup is due to selection rather than simply drift, since it was by far the strongest signal in the pcadapt or GEMMA-LMM analysis of the Jatt/Chowdhry subgroup (minimum p=3×10^−9^ in pcadapt), with no SNPs outside this region having p<10^−6^.

## Discussion

We have carried out the first large-scale investigation of population structure and demographic history of Pakistanis. We found genetic structure in the cohort reflecting the influences of the *biraderi* social stratification system, with some subgroups forming identifiable and homogeneous clusters (Figures 1 and 2). Our analyses suggest that these subgroups come from a common ancestral population but diverged from one another within the last 70 generations (1,500-2,000 years) (Figure 3a). This is consistent with an earlier finding that the transition from intermarrige to strict endogamy on the Indian subcontinent started from about 70 generations ago^74^, concurrent with or immediately after the drafting of the ancient Law Code of Manu that described a ranked social stratification system^31^.

Historically, intra-*biraderi* marriages and consanguinity were practised to solidify socio-economic bonds ^75,76^. *Intra*-biraderi marriages continue to be very common in the Bradford Pakistani community, constituting >90% of marriages in the Gujjar, Pathan, Jatt, and Bains, ≥80% in the Syed, Qasabi, Rajput, Awaan, and Choudhry, and ~60-63% in Qureshi, Malik, and Sheikh in the current BiB participants ^76^, according to the self-reported questionnaire data. Interestingly, the subgroups we inferred to be the most genetically homogeneous and with the highest IBD sharing (Figure 3a) are generally those with the highest self-reported rates of *intra*-biraderi marriage. Endogamous practices became stronger, particularly amongst the elite classes, during the Mughal Empire (mid-1500s to mid-1800s) ^30^, and then amongst all classes under the British Raj (mid-1800s to 1947) when the laws of land ownership changed ^30,31^. This could be the explanation for the decrease in effective population size seen in all subgroups starting 15-20 generations ago (~375-580 years ago if the average generation time were 25-29 years) (Figure 3b). However, there is considerable uncertainty in the average generation time; according to historical records, it was not uncommon for women in South Asian communities to marry and have children as early as 12-18 years old ^77,78^. Hence, we should be cautious about attributing these changes in N_e_ to demographic changes or historical events at particular time points.

To explore the elevated rates of homozygosity and IBD sharing in our cohort, we leveraged new theory (^62^ and Methods) to estimate the recent consanguinity rates in founder populations, while, for the first time, properly accounting for ROH attributed to historical endogamy. We found that for both the Jatt/Choudhry and Pathan subgroups, a naive estimate of spousal kinship based on self-reported parental relatedness underestimates the proportion of the genome in ROH segments, and thus, the true kinship coefficient between spouses must be higher. Stronger kinship would be consistent with the fact that 67% of the Pakistani mothers who reported being related to their partner also reported that their parents were related. A limitation of our approach was that we did not explicitly model this more complex pedigree structure. Another is that we have little power to identify historical changes in the consanguinity rate (Supplementary Figure 22). This is because, as we go back in time, the ROH and IBD distributions are similarly affected both by the average kinship between parents, *k*, and by *N_e_*, so we can only infer the product *N_e_* (1 – 3*k*).

Our results suggest that, even in the absence of close consanguinity, increased homozygosity due to endogamy is likely to be contributing to recessive disease burden ^79^ and the elevated frequency of rare homozygous knockouts ^12,80^ in this population. Offspring of unrelated parents from the same subgroup have on average 2.6-times more rare, putatively deleterious homozygous coding variants than offspring of unrelated parents from different subgroups (Supplementary Figure 23), which may be proportional to their risk of recessive disorders. However, we note there is a wide confidence interval on this estimate (i.e. [1.5-7.0]-times more).

We hypothesised that rare disease-causing variants might have increased in frequency in certain Pakistani subgroups through founder events. We did find a few dozen coding variants with significantly altered frequency amongst the three largest subgroups compared to the rest of the BiB Pakistani samples (Supplementary Table 15), but none of these had compelling evidence for pathogenicity. We identified multiple variants in the HLA region at significantly altered frequency in the Jatt/Choudry subgroup, which may be due to selection rather than drift. The HLA region has been identified as a target of selection in different human populations and is involved in immunity against pathogens but also implicated in other phenotypes such as high altitude adaptation ^73,81–83^. A larger sample size and a population-specific HLA reference panel would be required to fine-map the most differentiated variants in this region in the Jatt/Choudhry subgroup, which would be a prerequisite of forming hypotheses about which phenotypes might have been the target of selection.

This study has several limitations. Although we had samples from fifty-six distinct groups, many of them had a small sample size that would not allow reliable group-level inference about demographic history or founder events. The majority of our sample comprised individuals with Pathan, Punjabi or Kashmiri ancestry residing in the UK; additional data from the UK and Pakistan, including from other ethnic groups such as the Baloch and Sindhi, would allow us to explore how generalizable our findings are. Furthermore, our study was based primarily on SNP-genotype data, which, although highly valuable for describing population structure, did not allow the finer scale demographic inference that would be possible with whole-genome sequence data. For example, whole-genome sequence data on more males would allow us to leverage a larger number of Y chromosomal markers to further explore whether *biraderi* membership has indeed been passed down patrilineally in recent time.

We have presented the largest investigation to date of fine-scale population structure and history in Pakistanis. We have shown how the *biraderi* social stratification system has played a significant role in shaping the population structure in Pakistani communities over the last 70 generations. Endogamous practices have led to greatly elevated IBD sharing as well as increased homozygosity, which is likely to have implications for disease risk on top of the high rates of consanguinity. Larger sample sizes and data on Mendelian disease diagnoses (as opposed to proxy estimates of risk) will be needed to quantify the risk of recessive disorders for offspring of intra-versus *inter*-biraderi marriages accurately, noting that our results imply that this differs between *biraderi*, and that certain disorders may be enriched in particular *biraderi* as a result of founder effects. Furthermore, future studies of disease genetics in Pakistanis should consider our findings in order to best control for population stratification due to recent structure in genome-wide association studies ^16^ and potentially to increase power by exploiting the IBD within some groups as a proxy for rare variant sharing ^84^.

## Materials and methods

### Dataset

We had genetic data from 7,180 individuals of Pakistani ancestry and 6,818 of White British ancestry (<u>Supplementary Tables 1 and 2</u>) from the Born in Bradford (BiB) cohort. These were predominantly from mothers and their children, with some fathers also included. The children were only included for the analyses in Supplementary Figure 17, since we needed to restrict the population genetic analyses to unrelated individuals. The samples were genotyped using two chips: 1) the Infinium CoreExome-24 v1.1 BeadChip (~550K SNPs), and 2) the Infinium Global Screening Array-24 v1 (GSA; ~640K SNPs). We also analysed whole-exome sequencing (WES) data for 2,484 Pakistani individuals who were also genotyped. Recruited mothers were asked to fill in a self-reported questionnaire including, for the Pakistani mothers, questions about *biraderi* groups, places of birth and parental relatedness (consanguinity).

### Quality control of genotype chip data

Data quality control was performed using PLINK v1.90b4 ^85^. We required <1% missingness per SNP and <10% missingness per individual. Duplicated variants were removed. We removed 110 samples with sex discrepancies, 123 genetic duplicates, 465 individuals who were not genetically related to someone who should have been a first-degree relative, and five individuals who had an inferred first-degree relative who should not have been related. Genetic ancestry was assigned with principal component analysis (PCA) using the selfreported information on ethnicity, and an additional 52 samples were filtered out because their declared ethnicity was different from their genetic ancestry inferred using EIGENSOFT 7.2.1^37,86^. SNPs with Hardy-Weinberg Equilibrium p-value < 1×10^−6^ were removed considering Pakistani (3,348 CoreExome variants, 1,435 GSA variants) and White British (305 CoreExome variants, 224 GSA variants) separately. These filters resulted in a dataset of 476,816 autosomal SNPs (246 mtDNA SNPs) and 14,624 individuals on the Core Exome chip and 598,326 SNPs (583,667 autosomal, 1,056 Y chromosome) and 4,398 samples on the GSA. For this study, we focused primarily on individuals of Pakistani ancestry, but in some figures the White British individuals were used for comparison. Results presented are from the CoreExome chip data unless otherwise stated, since this gave a larger sample size, and taking the intersection of the two chips would have left insufficient SNPs for most analyses.

We performed the demographic inferences on mothers genotyped on the CoreExome chip using common autosomal SNPs (251,853 SNPs with minor allele frequency (MAF) > 0.01). For the PCA and ADMIXTURE analyses, we filtered out SNPs in high linkage disequilibrium (LD) (r^2^>0.5). Fathers from the GSA dataset were used for the Y chromosome analysis. This resulted in a dataset of 3,081 Pakistani and 2,873 white British mothers and 2,588 Pakistani children on the CoreExome chip and 601 mothers and 235 Pakistani fathers on the GSA.

### Inference and removal of relatives

A critical step in population genetic analysis is identifying and removing close relatives. In endogamous populations, distinguishing recent familial relatedness from population structure is challenging because both show genetic similarity through allele sharing ^87^. We inferred relatedness coefficients with KING, which has been reported to be robust to population structure ^36^, unlike PLINK’s 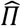 estimator. We compared the original kinship estimate from KING (called ‘KING-robust’ in ^36^) to a new estimator in KING, PropIBD, which integrates IBD segment information to infer sample relationships for 3rd and 4th degree relatives more accurately (http://people.virginia.edu/~wc9c/KING/manual.html). Comparisons of these different estimators on the mothers genotyped on the CoreExome showed that PropIBD identified more third-degree and closer relatives than the other methods (Supplementary Figure 2a,b): KING ProbIBD removed 881 samples, KING-kinship removed 741 samples, and PLINK’s 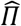 removed 821 samples.

To try to determine which of these estimators was more accurate, we compared the genetic estimates of kinship to self-reported relationships for 196 Pakistani mother-father pairs for whom the relationship had been declared by the mother on the questionnaire. For these, since mothers and fathers had been genotyped on different arrays, we took the intersection of CoreExome chip and GSA (134,218 SNPs with MAF>0.01) and ran KING on this (Supplementary Figure 2c,d). The results suggest that KING-kinship may be underestimating true kinship for third-degree relatives: 21/75 (28%) self-declared 1st cousins were called more distant than third degree relatives by KING-kinship, versus only 4/75 (5%) with KING-PropIBD. PropIBD called 38/75 (51%) self-declared 1st cousins as second-degree relatives. This seems to be mostly driven by inbreeding in the previous generation, since when we restrict to couples for whom both the mother’s and father’s parents were reported to be unrelated, KING-kinship and KING-PropIBD gave very concordant estimates (Supplementary Figure 2d).

To be conservative in our population genetic analyses, we decided to use the relatedness estimates from KING-PropIBD to exclude relatives. We removed one sample from each pair of individuals with third-degree relatedness or above (i.e. PropIBD>0.0884). We retained 2,200 Pakistani and 2,520 White British mothers and 1,616 Pakistani children on the Core Exome chip and 544 Pakistani mothers and 228 Pakistani fathers on the GSA.

### *Biraderi* categorisation

On the questionnaire, Pakistani mothers in BiB were asked to state their own *biraderi*, that of each of their parents, and that of their husband and of his parents. We cleaned the *biraderi* membership data, checking for spelling errors and combining groups with variable spellings of the same group. We determined that fifty-six distinct groups had been reported. To ensure we were assigning individuals to the *biraderi* that was most consistent with their parental *biraderi*, we compared the mother’s self-reported *biraderi* to the *biraderi* reported for each of her parents, and likewise for the father’s *biraderi*. The *biraderi* for a mother or father was set to missing if it was not reported (N=645), or if her/his parents’ *biraderi* was not reported (N=58) or was discordant with his/her own (N=118). The *biraderi* for a child was assigned following the same approach used for mothers and fathers. We assigned 2,324 mothers (1,652 unrelated), 1,568 children (906 unrelated) and 169 fathers (164 unrelated) unambiguously to *biraderi* groups.

### Population structure

For comparison with other modern worldwide populations, we merged our dataset with the Human Genome Diversity Project (HGDP)^88^ and 1000 Genomes Project Phase 3 ^21^. For comparison with modern and ancient individuals, we combined our samples with a dataset of published genotypes from modern and ancient individuals (https://reich.hms.harvard.edu/datasets) ^18^. In both cases, we used GRCh37.

Principal component analysis (PCA) was performed on the pruned datasets using EIGENSOFT 7.2.1 ^37,86^. For the worldwide dataset, the eigenvectors were computed using the non-BiB datasets and the individuals from BiB were projected onto them (Supplementary Figure 3a,b). For Supplementary Figure 3c, to ensure that the BiB Pakistanis were not dominating the structure due to their large sample size, we computed the PCA using HGDP Pakistani populations and just 25 BiB Pakistanis, then projected the remaining BiB Pakistanis onto them.

ADMIXTURE v1.3^38^ was run on the pruned datasets, and the cross validation (CV) error was calculated for identifying the best *K* value, which was found to be 4 for Figure 1c and 8 for Supplementary Figure 4a. We separated Rajput-A and Rajput-B according to the proportion of the red component in the ADMIXTURE analysis, defining Rajput-B individuals as those for which the red component made up >40% of their ancestry. Rajput-A individuals had a maximum red component proportion of 18%.

Genetic affinity with other worldwide populations was tested by computing *f*-statistics using ADMIXTOOLS v6.0. We computed both *f3* and *f4*-statistics. Outgroup *f3*-statistics were computed with the phylogeny *f3* (Bradford Pakistanis, X; Mbuti), where X represents another worldwide population. Standard errors were obtained using blocks of 500 SNPs. *f4*-statistics were computed using qpDstat with f4mode:YES and with the phylogeny *f4* (W, X; Y, Chimpanzee), where W represents the *biraderi* groups reporting Arabic ancestry (Qureshi, Sheikh or Syed), X are all the other Bradford sub-groups and Y represents Middle East populations^39^. Positive values indicate gene flow between W and Y. *f*-statistic tests with Z-score > 3 were considered significant.

Uniform Manifold Approximation and Projection (UMAP) was computed from the top 20 PCs with default parameters in the BiB dataset using the *uwot* R package ^89^. To compare genetic to geographic distance, we first assigned 547 mothers to a village of origin in Pakistan using either the mother’s own self-reported village of origin, or that of her parents if she was born in the UK and reported that her parents were born in the same Pakistani village. We then determined the latitude and longitude of the villages in order to define geographic distances between them. Genetic (measured as UMAP1 and UMAP2 vectors) and geographic distances were computed as Euclidean distances using the dist R function with method=Euclidean. Correlation between genetic and geographic distance was estimated using a Mantel test implemented in the *Ade4* R package (mantel.rtest function)^90^. The number of permutations used for the Mantel test was 9999.

We ran fineSTRUCTURE v4.0.1^54^ to infer fine-scale population structure in the BiB Pakistani mothers. The haplotypes were phased using SHAPEIT v2.12^91^ using the 1000 Genomes Phase 3 dataset as a reference panel. We generated the co-ancestry matrix using ChromoPainter and used it to run fineSTRUCTURE with 1,000,000 burn-in steps and 1,000,000 iterations. A PCA on the co-ancestry matrix using the *prcomp* R function (Supplementary Figure 7) confirmed the structure in the tree (Figure 2). We ran the algorithm twice on the dataset with the major subgroups to make sure the tree structure was robust (data not shown). We also ran it on the dataset including all subgroups to confirm that the clusters were consistent (Supplementary Figure 9).

Given the heterogeneity of the genetic clusters inferred in our cohort (Figure 2), we defined self-reported groups as homogeneous if at least 60% of the individuals from each selfreported subgroup fell in the same genetic cluster inferred by fineSTRUCTURE. We then estimated pairwise F_ST_ (calculated with the Weir and Cockerham’s method using the program 4P^92,93^) between homogeneous self-reported groups within a fineSTRUCTURE cluster, and combined two groups if they had F_ST_< 0.001 (5th percentile of the empirical distribution for all pairs of groups shown in Supplementary Figure 8b). For demographic analyses, we included only the homogeneous self-reported groups that had at least 20 individuals. These were: Awaan+Syed from Cluster 2, Bains+Rajput-B from Cluster 9, Jatt+Choudhry from Cluster10, Arain from Cluster 4, Gujjar from Cluster 5, Kashmiri from Cluster 1, Pathan from Cluster 8 and Qasabi from Cluster 7. Sample sizes for these groups are shown in <u>Supplementary Table 7</u>.

Chromosome Y haplogroups were defined with yhaplo ^94^ and mtDNA haplogroups with Haplogrep2 ^95^. A median joining haplotype network for the Y chromosome data from BiB Pakistani fathers was constructed with PopART v1.7 ^96^.

### Divergence time estimation

We calculated the time of divergence between the homogeneous sub-populations and genetic clusters using the approach described in McEvoy *et al*.^97^ and implemented in the NeON R package ^55^. The 95% confidence intervals for the times of divergence were calculated using a jackknife procedure, leaving out one chromosome each time.

### IBD segment calling

We conducted several analyses using identical-by-descent (IBD) segments between the Pakistani individuals, called using IBDseq v.r1206 ^98^ and/or GERMLINE 1.5.3 ^99^. IBDseq requires unphased data whereas GERMLINE phased haplotypes. IBDseq ran with default parameters. For GERMLINE, we phased the data with Eagle v2.4.1 ^100^ using 1000 Genomes Project Phase 3 genetic maps. IBD segments were defined using GERMLINE with the parameters -bits 75 -err_hom 0 -err_het 0 -min_m 3 -h_extend, and the HaploScore algorithm ^101^ was used to remove false-positive IBD tracts (genotype error=0.0075, switch-error=0.003, mean overlap = 0.8).

In plotting the unfiltered IBD calls along the genome, we noticed that a suspiciously high number of IBD calls were being made in particular regions (Supplementary Figure 24), and that these regions were enriched for gaps or overlapping centromeric regions, suggesting that these were artefacts. We thus excluded IBD segments overlapping these regions: chr1:120-152Mb, chr2:88-102Mb; chr9:38-72Mb; chr10:17-19Mb and chr11:0-3Mb (Supplementary Figure 25; “filter 1” in Supplementary Figure 26). We furthermore excluded a small number of outlier IBD segments longer than 23cM that were seen with identical coordinates in more than ten pairs of unrelated (PropIBD<0.084) individuals, and some IBD segments >25cM partially overlapping the centromeric region of chr15, which were seen with identical coordinates in more than five pairs of unrelated individuals (“filter 2” in Supplementary Figure 26). Finally, we excluded IBD segments overlapping the HLA region and centromeres.

### Estimating historical effective population size

We estimated historical changes in effective population size (N_e_) for both Pakistani subgroups and White British individuals using IBDNe v.19Sep19.268 ^60^. As recommended, we ran IBDseq ^98^ to identify IBD segments with default parameters. We then excluded IBD chunks overlapping problematic regions as explained above, although in practice we found this had a minimal effect on the results. We ran IBDNe with the default parameters, except that we set a lower limit of 5cM for the IBD segments considered. We also ran IBDNe using the IBD calls from GERMLINE and found that this gave systematically higher N_e_ estimates than IBDseq, although the trajectories were very similar, particularly for the larger groups (Supplementary Figure 27).

### Calculating IBD scores

We computed the IBD score developed in Nakatsuka *et al*. ^34^ from GERMLINE IBD calls to quantify the extent of founder events in each population using IBD segments. Nakatsuka *et al*. had used IBD segments >30cM to define and exclude possible close relatives (on top of a filter based on PLINK’s 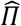), and then considered the total length of segments between 3 and 20cM. The appropriateness of these cutoffs depends on the patterns of IBD sharing within a population of interest, as well as the density of SNPs being used to call IBD segments. We found that in the BiB Pakistanis, a nontrivial fraction of individuals who were estimated to be unrelated by KING had an IBD segment between 20 and 30cM (Supplementary Figure 13), and due to the sparsity of SNPs on the CoreExome chip, we suspected IBD segments <5cM were not reliably estimated. Hence, we selected slightly different filters when preparing Figure 4. Specifically, in addition to the aforementioned removal of individuals who were third-degree relatives or closer based on the KING PropIBD metric (PropIBD<0.0884), we also removed one individual from each pair of samples sharing an IBD chunk > 40 cM, to be sure that we were not mistakenly including closely related individuals.

We calculated IBD scores as the total length of IBD segments between 5 and 30 cM detected between individuals in the same subgroup divided by the total number of possible pairs i.e. 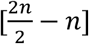, where *n* is the number of samples in that subgroup. We then standardised each IBD score by the score for the Finnish individuals from the 1000 Genomes Phase 3 ^34^. Standard errors for IBD scores were calculated using a weighted block jackknife for each chromosome, and 95% confidence intervals were defined as the IBD score ± 1.96 × standard error. We also computed IBD scores using the same set of filters used in Nakatsuka *et al*. (IBD segment filter > 30 cM to identify relatives, then counting IBD segments between 3 and 20 cM) and found that the results were similar (Supplementary Figure 14).

### Inferring gene flow between Pakistani subgroups

Gene flow was inferred using the Treemix v1.13 approach ^56^ and *f3*-statistics ^19,39^. We ran the Treemix analysis with default parameters, without the -root option. We added migration edges (-m) until we reached a total variance explained of 99.5%. The *f3* statistics were calculated in the form of *f3* (target, source 1, source 2) using ADMIXTOOLS v6.0, and provide evidence that the target population is derived from an admixture of populations related to source 1 and source 2. We tested all possible combinations of target and source populations in our dataset. Standard errors were obtained using blocks of 500 SNPs. Tests with a Z-score < −3 were considered significant.

### ROH calling

We used three different ROH callers: bcftools/roh, GARLIC v1.6.0a, and PLINK. The results in Figures 4 and 5 and Supplementary Figures 16, 17, and 18 are based on bcftools/roh calls, and in Supplementary Figure 20 and Supplementary Figure 23 we compare all three callers.

With bcftools/roh, we used the -G 30 flag. We noticed an excess of apparently artefactual ROHs spanning long gaps between SNPs around centromeres, so we removed ROHs < 10Mb overlapping centromeres. In practice, this made very little difference to the ROH footprint (Supplementary Figure 20). For GARLIC ^63,102^, we set the number of resamples for estimating allele frequencies (--resamples) to 20 and we assumed a genotyping error rate (-error) of 0.001. We used the --auto-winsize flag to guess the best window size based on the SNP density. For PLINK, we followed previous publications ^103,104^ and used the following parameters: --homozyg-window-snp 50 --homozyg-snp 50 --homozyg-kb 1500 --homozyg-gap 1000, --homozyg-density 50 --homozyg-window-missing 5 --homozyg-window-het 1.

### Analysis of F_ROH_

F_ROH_ was determined for each individual by summing the lengths of the (autosomal) ROHs in base pairs and dividing by the length of the autosomal genome. We fitted an ANOVA to assess the effect of self-reported consanguinity and self-reported subgroup and their interaction on F_ROH_ in the mothers. We calculated the partial r^2^ of each factor using the etasq() function in R. The categories of self-reported consanguinity are shown in Supplementary Table 13; for this analysis, we combined ‘other blood’ and ‘other marriage’ into a single category. The figures given in the Results section come from bcftools/roh calls, but very similar results were obtained with PLINK and GARLIC (data not shown). We also calculated F_ROH_ in the BiB children from the bcftools/roh calls and stratified these estimates according to the parents’ birthplace: both parents born in the UK, one parent born in the UK and one in Pakistan, and both parents born in Pakistan.

### Analysis of ROH and IBD distributions

We applied theory from ^62,105^ to determine the expected length distribution and genomic footprint of ROHs and IBD segments given a particular historical N_e_ trajectory and rate of consanguinity. In the model developed in the above papers, there are *N_e_* pairs of parents, who, in each generation, have probability *r*_1_ of being (full) siblings, *r*_2_ of being (full) first cousins, *r*_3_ of being second cousins, etc., up to a certain degree *n*. A key parameter of the model is the average kinship between parents, 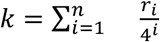, which is the probability of a random chromosome in each parent coalescing due to consanguinity. It is assumed that the parents are related only through a single path of degree 1, …, *n*, or are otherwise unrelated.

In ^62^, a Markov chain was derived for the evolution of the state of two chromosomes. The possible states are as follows: the chromosomes are in unrelated individuals, in one of each parent, in the same individual as two homologous chromosomes, or in the same individual as a single chromosome (coalescence). We then considered *t_between_*, the time to the most recent common ancestor (TMRCA, or coalescence time) of two chromosomes sampled from two unrelated individuals, and *t_within_*, the TMRCA of the two chromosomes of an individual. In refs. ^62,105^, the mean and variance of *t_between_* and *t_within_* were computed using a first-step analysis. An approximation of the distribution was derived using a separation-of-time-scales analysis, The results showed that *t_between_* is approximately exponential with rate 1/[4*N_e_* (1 – 3*k*)]. [The factor of 4 is because there are *N_e_* pairs of parents, thus 2*N_e_* individuals, and 4*N_e_* chromosomes.] *t_within_* has probability *k*/(1 – 3*k*) of being O(1), i.e., representing a rapid coalescence within the family due to consanguineous unions, and is otherwise approximately exponential as *t_between_*.

Here, we assume that *t_between_* is distributed as in the standard coalescent, but with a population size trajectory scaled according to the theory as *N_e_*(*t*)(1 – 3*k*). Thus, in discrete time, it has the distribution 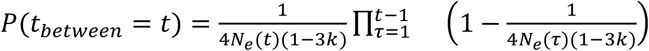. For *t_within_*, as the approximate distribution groups together all rapid coalescence events, we used a composite approach. For *t* ≤ 50, the distribution was computed numerically by running the exact Markov chain for 100,000 iterations. For *t* > 50, the distribution was set to 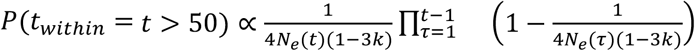 (as for *t_between_*), where the coefficient of proportion was set such that the entire distribution was normalized. Running the exact model for only *t* < 10 or *t* < 40 gave almost indistinguishable results.

We defined the genomic “footprint” of IBD and ROH segments as the proportion of the genome (in genetic map units) found in segments (of each type, respectively) of length between [*l*_1_, *l*_2_] ^106^. Given the distributions *P*(*t_between_*) and *P*(*t_within_*), we computed the footprint as follows. In ^107^, Ringbauer *et al*. (eq. (4) therein) showed that, in a chromosome of length *L*, the mean number of segments with TMRCA *t* of length in the small interval [*l,l* + Δ*l*] (in Morgan) is:

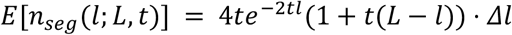

The mean chromosome length covered by segments (with TMRCA *t*) of length in [*l*_1_, *l*_2_], denoted *c*(*l*_1_, *l*_2_; *L*, *t*), is:

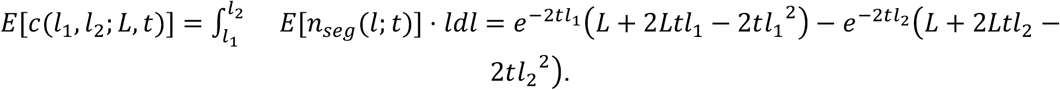

The mean length covered by segments of all TMRCAs is obtained by numerically summing over all TMRCAs, using their distributions (*t_between_* for IBD segments, who are between unrelated individuals, and *t_within_* for ROH segments):

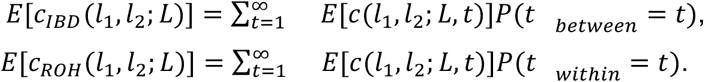

Finally, the footprint is obtained by summing over all chromosomes and dividing by the total genome length,

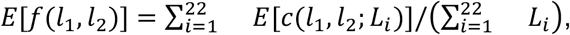

where *L_i_* is the length of chromosome *i*, in Morgans. The code implementing this model can be accessed at (https://github.com/scarmi/ibd_roh).

To generate the expected statistics shown in Figure 5 and Supplementary Figures 19, 20, and 22, we set *r*_1_ = 0 and estimated *r*_2_ and *r*_3_ using the fraction of mothers who declared that their parents were first cousins/first cousins once removed or second cousins, respectively. We set *r*_4_ to be the proportion of mothers who declared that their parents were related but that the relationship type was unknown or was described as ‘other blood’ or ‘other marriage’. This gave an estimate for *r* = (*r*_1_, *r*_2_, *r*_3_, *r*_4_) of (0, 0.235, 0.052, 0.155) for Pathan in cluster 8 and of (0, 0.425, 0.057, 0.151) for Jatt/Choudhry in cluster 10, giving naive kinship estimates 0.016 and 0.028, respectively. We also tried scaling the *r* vectors by a constant *c* to obtain higher kinship estimates for plugging into the model, and then compared the observed ROH footprint to the expectations given these values (Figure 5, Supplementary Figure 20). [Scaling *r* by *c* > 1 implies that the sum of its components may in some cases be greater than 1, which could represent multiple paths leading to the same relationship. The model still holds as long as the kinship satisfies *k* < 1/4.]

To investigate the effect of a sudden change in consanguinity rates at a certain point in the past, we tried reducing the kinship value to 1×10^−4^ at the time the distribution switched to the approximate model (*t* = 50). However, this proved to have only very subtle effects on the expected ROH and IBD segment footprint (Supplementary Figure 22), so we could not evaluate this possibility with the observed data.

For the N_e_ trajectory, we used a constant value of N_e_ for Supplementary Figure 19. For the other analyses, we used trajectory estimated with IBDNe for the last 50 generations (using the point estimate for Figure 5 and the bounds of the 95% confidence interval for Supplementary Figure 20), followed by a constant value for *t* > 50 (the IBDNe estimate at *t* = 50). [We divided the IBDNe population sizes by 2 before plugging them into the model, as the model assumes *N_e_* is the number of mating pairs.] We note that by using IBD data alone, IBDNe estimates *N_e_*(*t*)(1 – 3*k*) rather than *N_e_*(*t*). Thus, given the IBDNe inferred trajectory 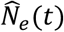, we set 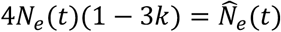 in the equations above.

To compute the ROH footprint in the real data, we averaged the total lengths of ROH segments (within each length interval) over all individuals in the group, and divided by the number of individuals. For the IBD footprint, we averaged the total lengths of IBD segments over all pairs of individuals in the group, and divided by 2*n*(2*n* – 2)/2 (where *n* is the number of individuals), which is the number of chromosome pairs in different individuals.

We restricted the real data analysis to ROHs and IBD segments greater than 5cM, since we suspected that segments shorter than this could not be called reliably with the CoreExome chip data, and less than 30cM, since there were few ROHs longer than this so the average footprint became very noisy. We used 1cM segment length intervals for both ROH and IBD, where each data point was plotted at the beginning of each interval. The empirical IBD footprint was plotted using the IBDseq calls filtered as described above, since these calls were used as input for IBDNe.

### Quality control of exome sequence data

Whole-exome sequence (WES) data were generated, and mapping and variant calling carried out as previously described ^12^ using BWA-MEM ^108^ and GATK ^109^ respectively. 2,311 samples were generated as part of this previous publication and had a median on-target coverage of 38X, and 473 additional samples were sequenced later to a median coverage of 50X. This gave a dataset of 2,784 individuals, of which 2,484 were Pakistani. We excluded variants that failed these criteria, based on standard GATK annotations:

- SNPs: QD < 2, FS > 60, MQ < 40, MQRankSum < −12.5, ReadPosRankSum < −8, or Hardy-Weinberg equilibrium p-value < 1×10^−6^.
- Indels: QD < 2, FS > 200, ReadPosRankSum < −20, or Hardy-Weinberg Equilibrium p-value < 1×10^−6^.

We set genotypes to missing if they had genotype quality (GQ) < 20, allelic depth p-value < 0.001, or depth < 7, then removed variants with more than 20% missingness. This resulted in a dataset of 1,931,299 variants (1,895,447 SNPs and 35,852 indels). The variants were annotated with the Variant Effect Predictor (version 95) using the LOFTEE plugin.

### Burden of rare damaging variants by parental relatedness level

For Supplementary Figure 23, we defined groups of individuals to compare based on the self-reported parental relatedness and self-reported parental subgroups. We extracted variants with minor allele frequency < 1% in the BiB Pakistani mothers from the exome sequence data. We then restricted to those whose most damaging annotation for any transcript was either:

- loss-of-function (LOFTEE high confidence), or
- missense and predicted to be “deleterious” by SIFT and “probably damaging” or “possibly damaging” by PolyPhen

We then simply counted the number of homozygous genotypes at these variants per individual (distributions shown in Supplementary Figure 23A), and calculated the ratio of mean numbers per group for Supplementary Figure 23B. The confidence intervals were calculated by bootstrapping individuals 1000 times within each of these groups.

### Inference of founder effects on specific variants

We tested for potential founder effects in particular subgroups using three approaches: 1) a PCA approach for outlier detection implemented in the pcadapt R package ^65^ applied to the CoreExome chip data; 2) a genome-wide association study-like approach implemented in GEMMA-LMM ^66^ applied to the CoreExome chip data; 3) a simple Fisher’s exact test on variants ascertained with whole-exome sequencing data, following the example in ^67^.

We compared allele frequencies in either Bains/Rajput-B in Cluster 9 (CoreExome: N=98; WES: N=64), Jatt/Choudhry in Cluster 10 (CoreExome: N=299; WES: N=195) or Pathan in Cluster 8 (CoreExome: n=213, WES: n=88) (“target group”), with a “control group” comprising individuals in Clusters 1-5 and 7 (CoreExome: N=723; WES: N=432) (Figure 2). We excluded Cluster 6 from the “control group” since it contained Bains/Rajput-B individuals who might be admixed. For both pcadapt and GEMMA-LMM, we included only polymorphic SNPs with an allele count (AC) ≥ 5 in the target+control groups to avoid false positives due to errors in assessing rare variants with SNP genotype data ^110^. For the Fisher’s exact test using whole-exome sequencing data we included variants with an AC ≥ 2 in the target+control groups.

For the pcadapt analysis, we first performed PCA on the genotype matrix generated with the function read.pcadapt from the PLINK files to verify that it was indeed PC1 that was separating the target from the control group. We then performed a component-wise genome scan according to the package instructions (https://bcm-uga.github.io/pcadapt/articles/pcadapt.html), specifying K=1 and method=componentwise. The method effectively computes the correlation between each SNP and the *k*th principal component of interest (in our case, PC1), and determines an associated p-value.

We carried out association tests with a univariate linear mixed model approach implemented in GEMMA-LMM, comparing the target to the control group. Firstly, we calculated a genomic relatedness matrix (GRM) with the -gk flag. We then conducted the association analysis (-lmm flag) using the GRM and the first 20 PCs as covariates.

We performed a two-sided Fisher’s exact test using the *fisher.test* function in R on variants ascertained with whole-exome sequencing data and on those that were significant with the pcadapt or GEMMA-LMM analysis on the CoreExome chip. We used the number of reference and alternate alleles for each variant in our target and control groups. Supplementary Table 14 includes the log-odds ratio calculated using the Haldane–Anscombe correction for comparisons with a zero-cell in the contingency table:

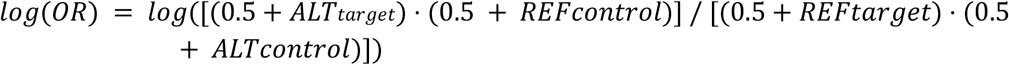

For all three tests we applied Bonferroni correction and considered SNPs with p-value < 1×10^−7^ as significant for the pcadapt and GEMMA-LMM analyses (0.05/N, where is the number of SNPs used in the analysis: Jatt/Choudhry=256,523, Bains/Rajput-B=256,603, Pathan=257,020) and p-value < 1×10^−7^ as significant for the Fisher’s exact test analysis on the whole-exome sequencing data (0.05/N, with N being: Jatt/Choudhry=279,643, Bains/Rajput-B=261,563, Pathan=263,107). We annotated variants using Ensembl Variant Effect Predictor (GRCh37) and generated Manhattan plots using the *qqman* R package^111^.

To determine the number of independent significant loci, we merged the CoreExome and exome-sequence data for each target group separately (Bains/Rajput-B, Jatt/Chowhury, Pathan), then calculated LD between significant variants. Variants with an r^2^> 0.2 were considered part of the same locus.

## Data availability

The genetic data and questionnaire data (covering self-reported consanguinity, village of origin and *biraderi* group) from Born in Bradford can be obtained as described here https://borninbradford.nhs.uk/research/how-to-access-data/.

## Acknowledgements

The study is only possible because of the enthusiasm and support of the children and parents recruited and the health professionals and researchers who make Born in Bradford happen, so we thank those individuals. Additionally, we thank Shahid Islam from the Bradford Institute for Health Research, two local British Pakistani leaders in Bradford, Mr. Ishtiaq Ahmed and Mr. Nazir Tabassum, and James Caron from the London School of Economics, for their insights into the historical and ethnographic background of the *biraderi* groups. We are grateful to Nathan Nakatsuka and David Reich for the IBD scores code, to Alan Bittles and Noah Rosenberg for their comments on the manuscript, and to Alissa Severson and Noah Rosenberg for assisting with the ROH analysis. S.C. was supported by the United States-Israel Binational Science Foundation grant 2017024.

## Author contributions

E.A., M.M., T.T. and H.C.M. analysed the data. K.E.H., Q.H., D.M., D.A.v.H. and M.M.I. carried out quality control on the data. S.A.D. and N.S. provided input on the *biraderi* ethnography and history. J.W. provided the data. S.C. contributed code and intellectual input on the IBD and ROH analyses. M.M.I. and H.C.M. supervised the study. E.A. and H.C.M. wrote the manuscript, with particular input from S.C., M.M., S.A.D. and N.S. All authors commented on the final manuscript.

## Supplementary Figures

**Supplementary Figure 1:**
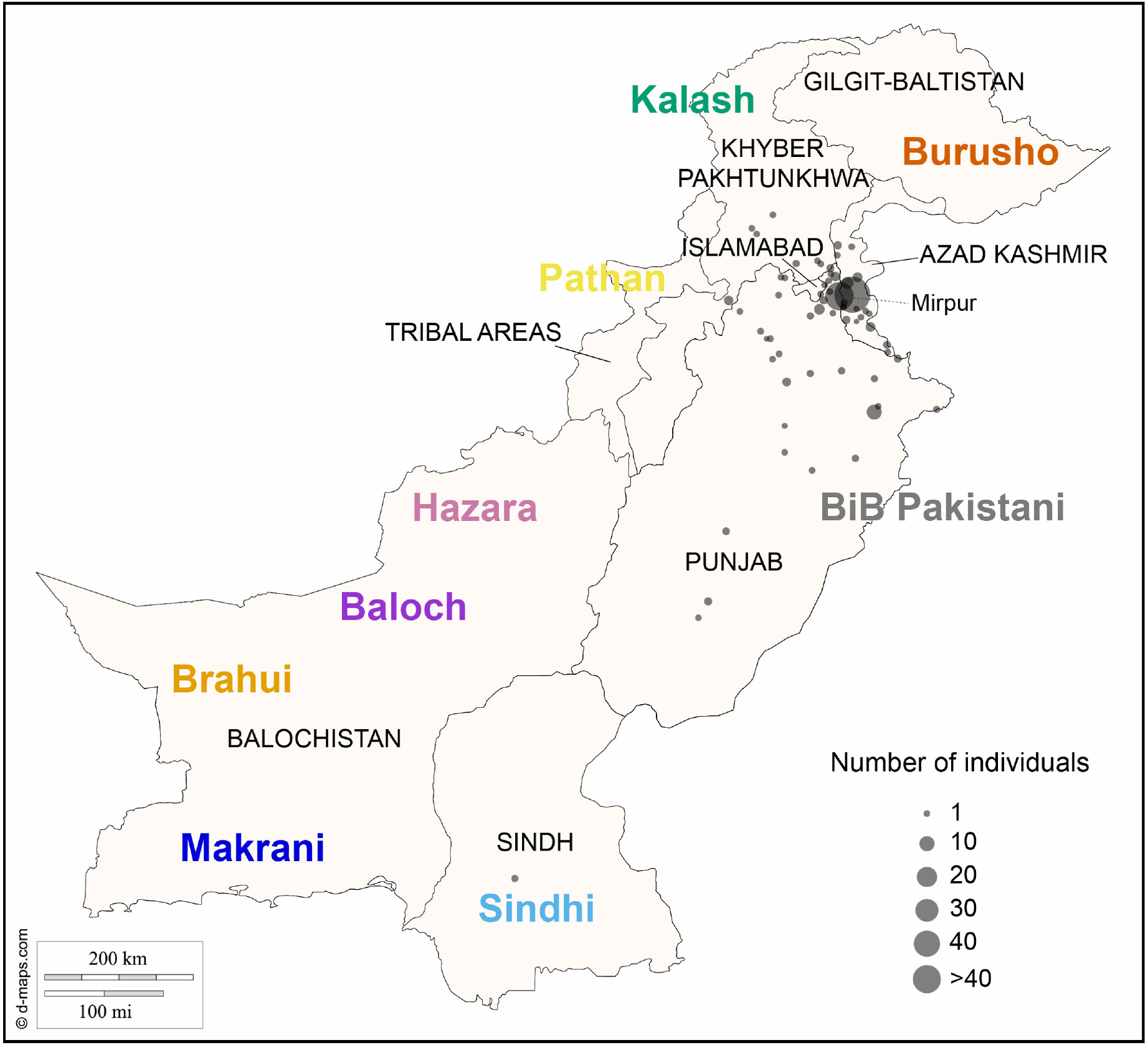
Map of Pakistan indicating the approximate origin of each of the HGDP populations in the coloured text, as well as the self-reported geographic origin location of the BiB Pakistani (see Methods). The size of the circles is proportional to the number of individuals reporting that they originate from that location. Most of the BiB Pakistani are from Mirpur in Azad Kashmir and northern Punjab. Note that although the Pathan are concentrated in the northwest of the country, many Pathan also live in other parts of Pakistan including Punjab, Sindh and Kashmir. The map does not include the boundaries of the disputed territory of Jammu & Kashmir.

**Supplementary Figure 2:**
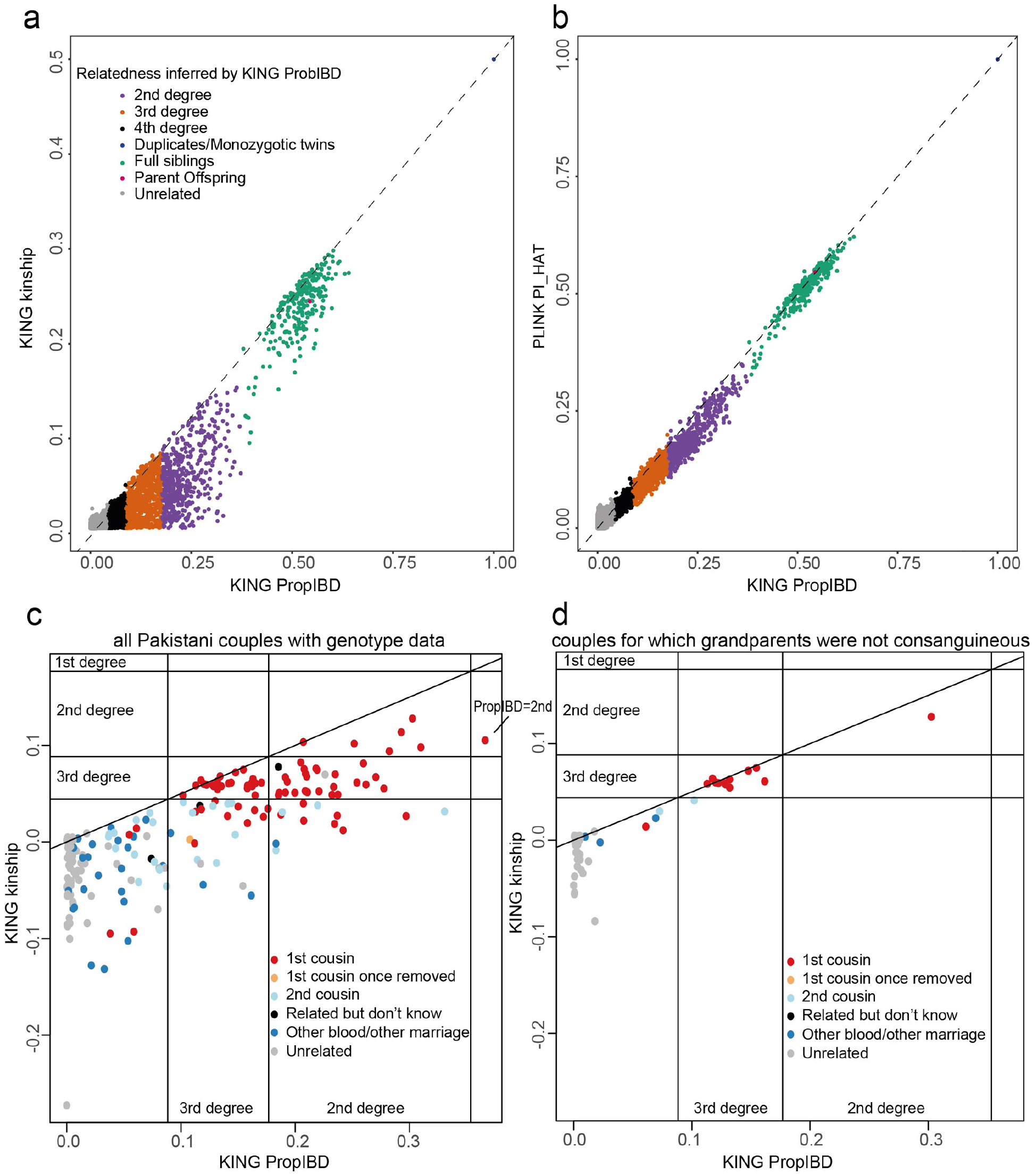
Relatedness checks. a-b) Relatedness estimates between all BiB CoreExome Pakistani mothers compared between three different metrics, where the colours indicate the relationship type inferred by KING-PropIBD. a) KING-kinship versus KING-PropIBD; b) KING PropIBD versus PLINK’s 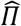 (PI_HAT). In (c) and (d), we compare the two KING estimators for the 196 Pakistani couples from the GSA/CoreExome intersection for whom we had self-declared relationship information. (c) shows all couples and (d) shows only those for which both the mother’s parents and father’s parents were reported to be unrelated. The vertical and horizontal lines indicate the recommended cutoffs for defining relatives, and the sloped line indicates the expectation if the two methods were performing identically.

**Supplementary Figure 3.**
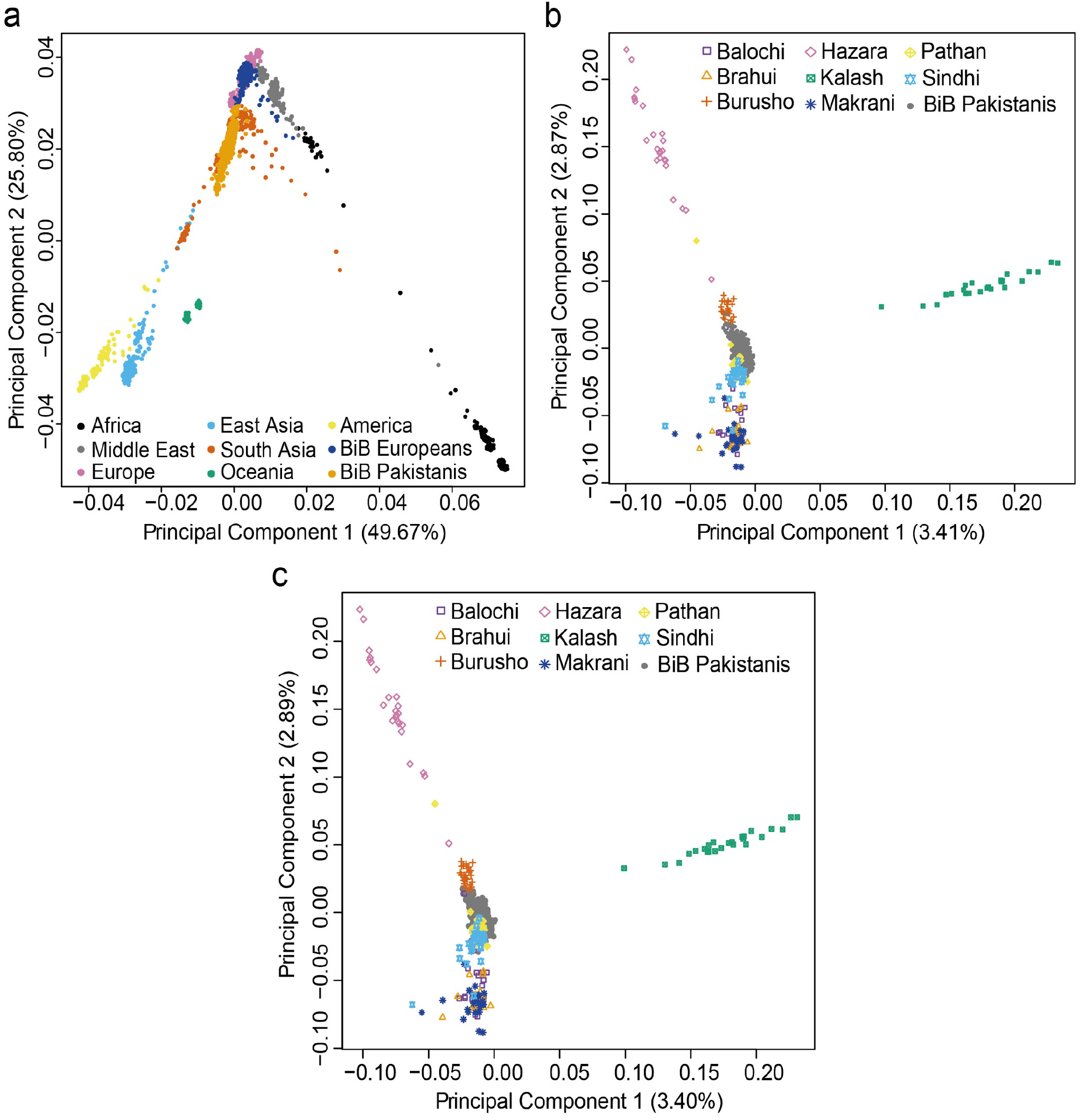
PCA of Born in Bradford mothers along with external datasets. a) PCA of the HGDP dataset with BiB Pakistani and European samples projected onto it, illustrating their position in a worldwide context. b) PCA of HGDP Pakistani samples, with BiB Pakistani samples projected onto them. c) PCA computed using HGDP Pakistani samples and 25 BiB Pakistanis, with the remaining BiB Pakistanis projected onto them. In both (b) and (c) the BiB Pakistanis cluster together on the PCA, on top of the HGDP Pathan and between the Sindhi and Burusho

**Supplementary Figure 4.**
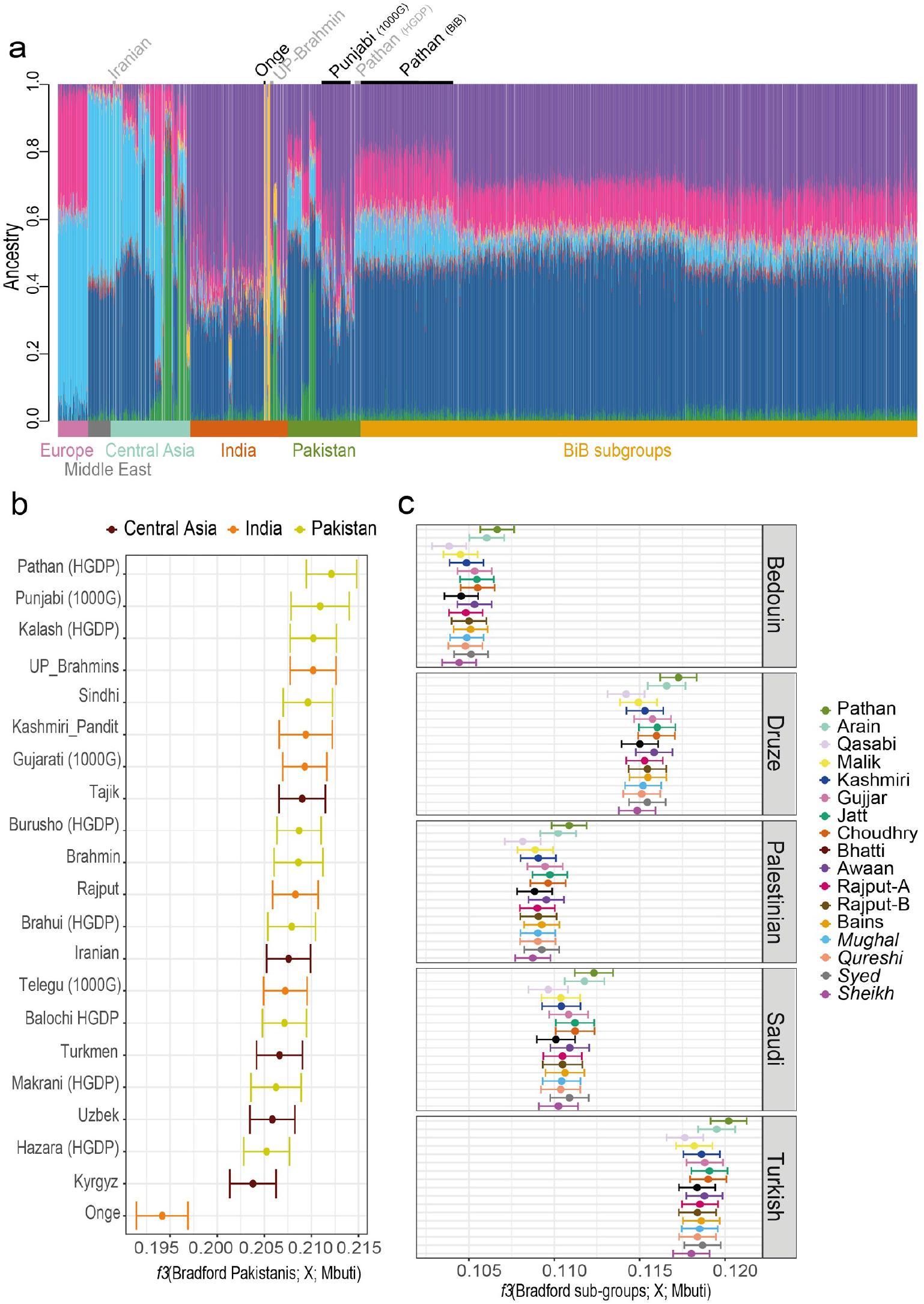
Genetic similarity between Bradford Pakistanis and other worldwide populations. a) Admixture plot (K=8) illustrating different ancestral components making up the various worldwide populations. BiB Pathans stand out compared to other subgroups and have a similar genetic profile to the HGDP Pathans. b,c) Outgroup *f3*-statistics of BiB Pakistanis compared to other worldwide populations. The x-axis represents the *f3*-statistics, computed with the phylogeny *f3*(Bradford Pakistanis, *X*; Mbuti), where *X* represents the indicated worldwide population. The higher the value, the higher the genetic sharing between the pair of populations tested. Error bars indicate standard errors. b) All BiB Pakistanis compared to other South and Central Asian groups. c) Self-reported subgroups compared to Middle Eastern populations indicated on the right. In *italics* are the self-reported groups that claim Arabic ancestry: Qureshi, Syed, and Sheikh.

**Supplementary Figure 5.**
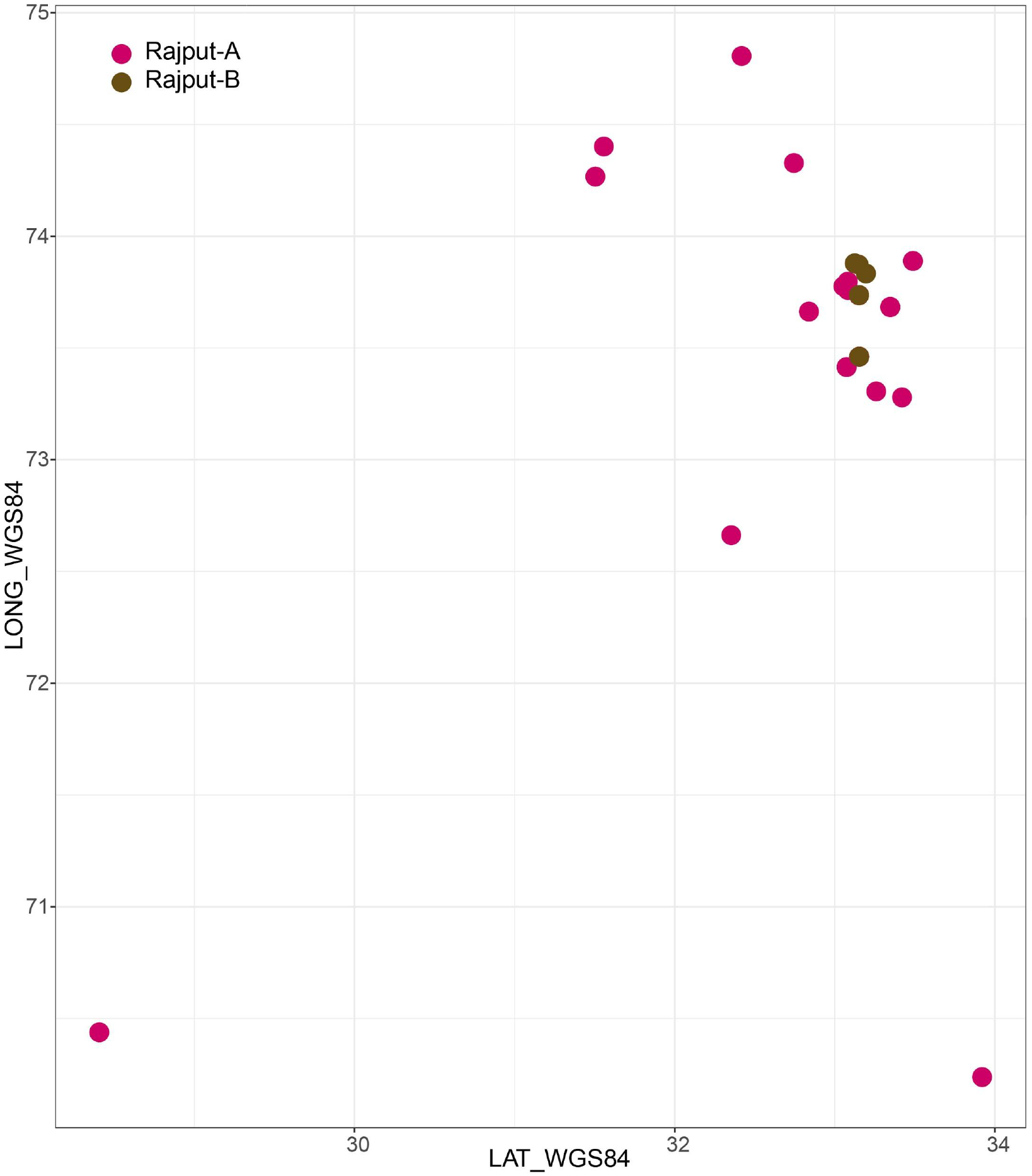
Comparison of the geographic origin of Rajput-A and Rajput-B individuals using the World Geodetic System (WGS-84) latitude and longitude coordinates. This is based on self-reported information about the mother’s own village of origin, or that of her parents.

**Supplementary Figure 6.**
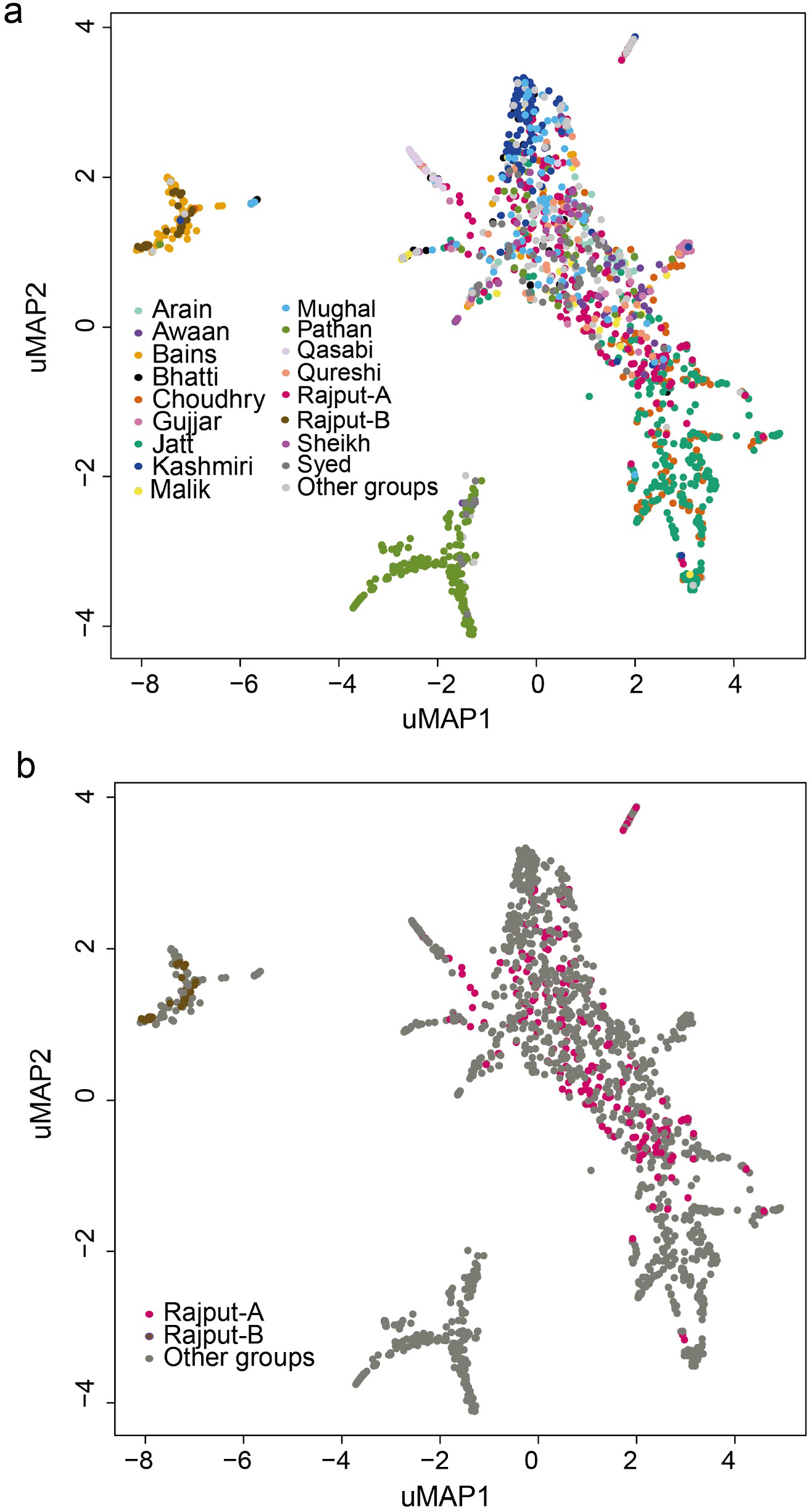
Population structure of the Bradford Pakistani subgroups as inferred with UMAP analysis using 20 PCs. a) UMAP plot coloured by major self-reported groups. b) The same UMAP plot but highlighting Rajput-A and Rajput-B samples in pink and brown respectively.

**Supplementary Figure 7.**
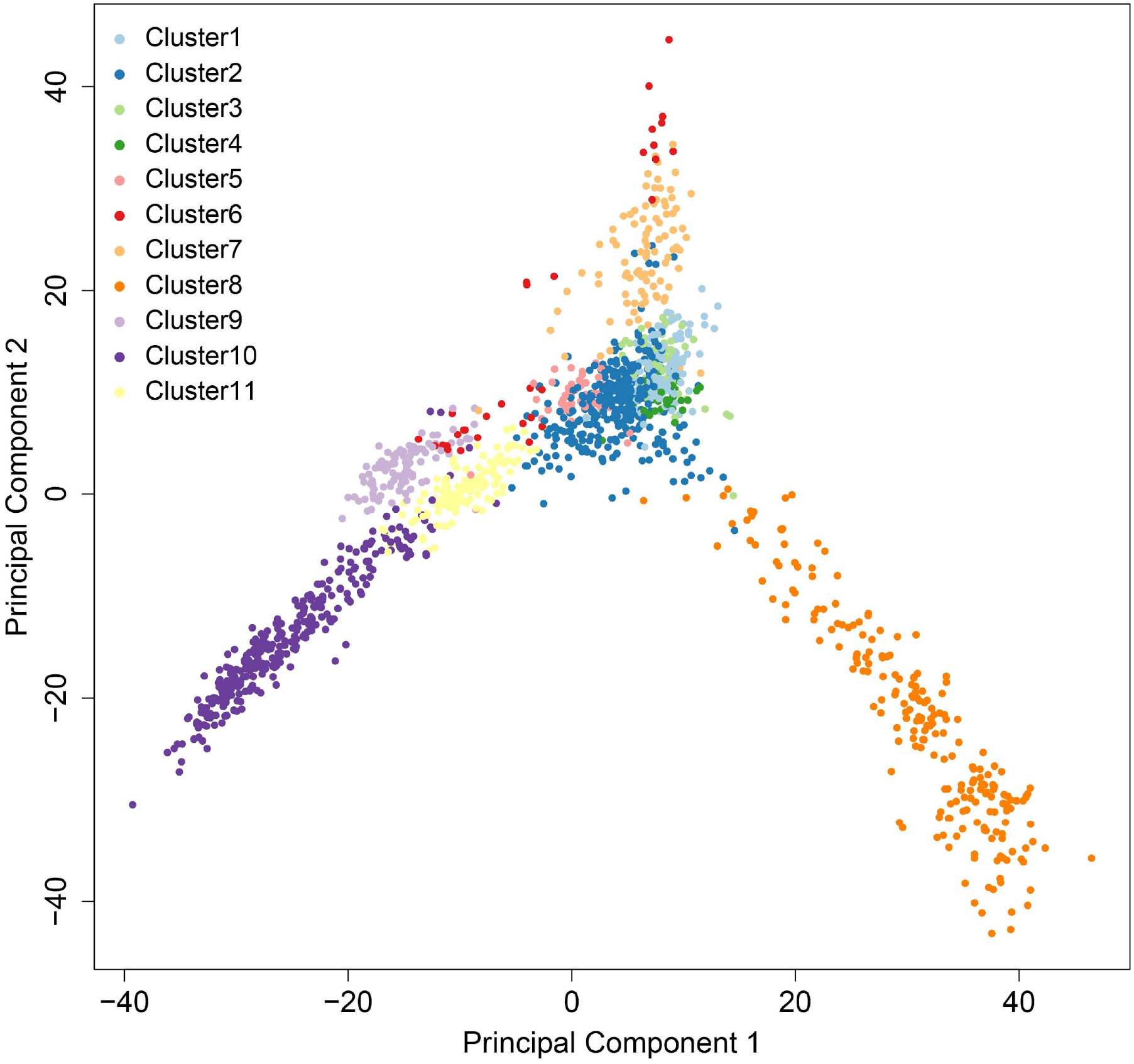
PCA on ChromoPainter co-ancestry matrix coloured by the genetic clusters defined by fineSTRUCTURE in Figure 2.

**Supplementary Figure 8.**
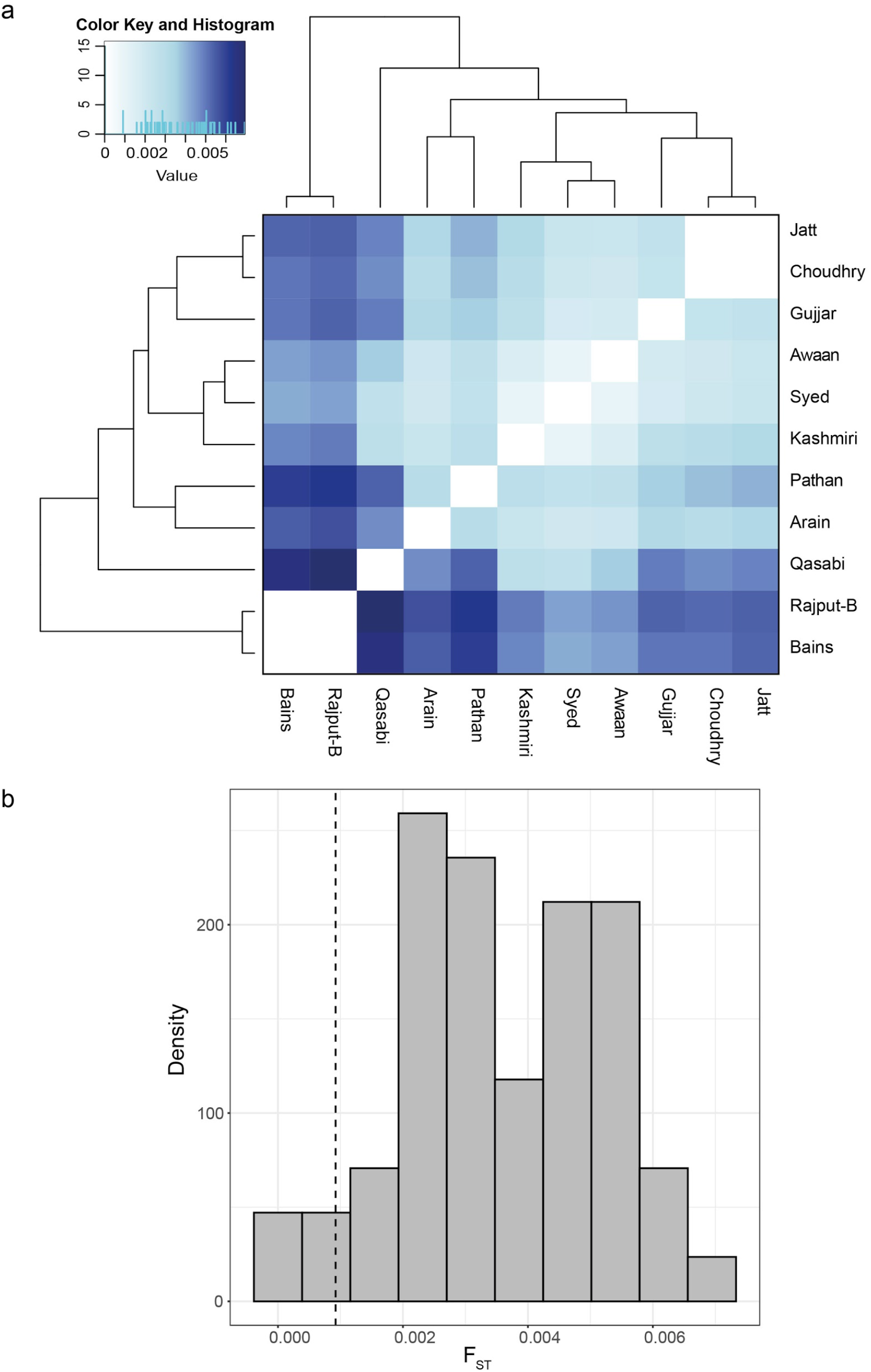
a) Heatmap of pairwise F_ST_ values for the subgroups, using only individuals who fell within the dominant cluster for that subgroup on fineSTRUCTURE (see Methods). b) Distribution of all pairwise F_ST_ values, with the dotted line denoting the 5th percentile of the empirical distribution.

**Supplementary Figure 9.**
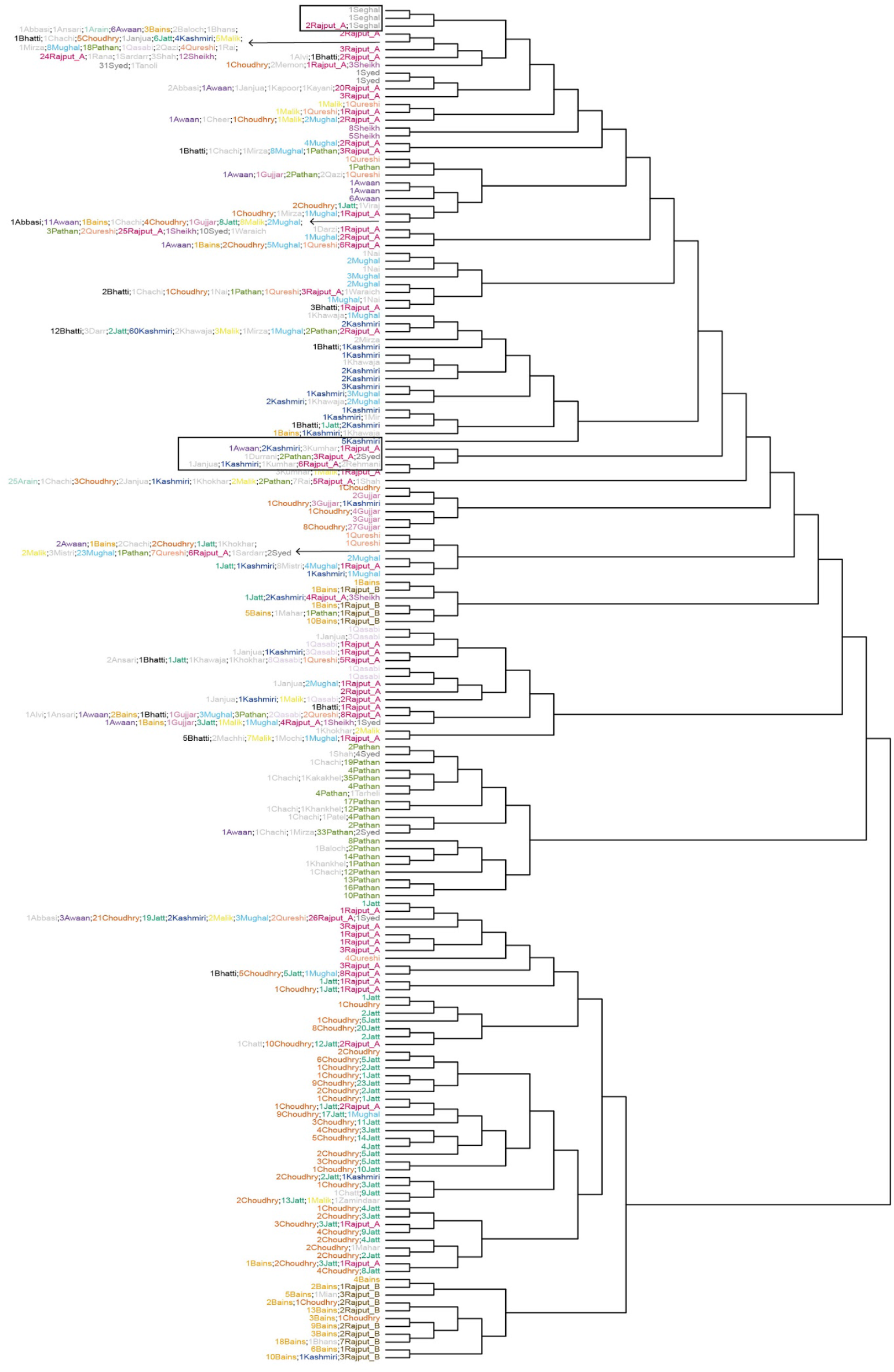
fineSTRUCTURE tree for all subgroups among Bradford Pakistanis. The tree illustrates the results of hierarchical clustering of the co-ancestry matrix defined using patterns of haplotype sharing from ChromoPainter. Each ‘leaf’ on the tree contains multiple individuals indicated by the number before the subgroup name. Boxed in black are additional individuals from other subgroups clustering together that were not included in Figure 2 (Kumhar, Rehmani, Sehgal).

**Supplementary Figure 10.**
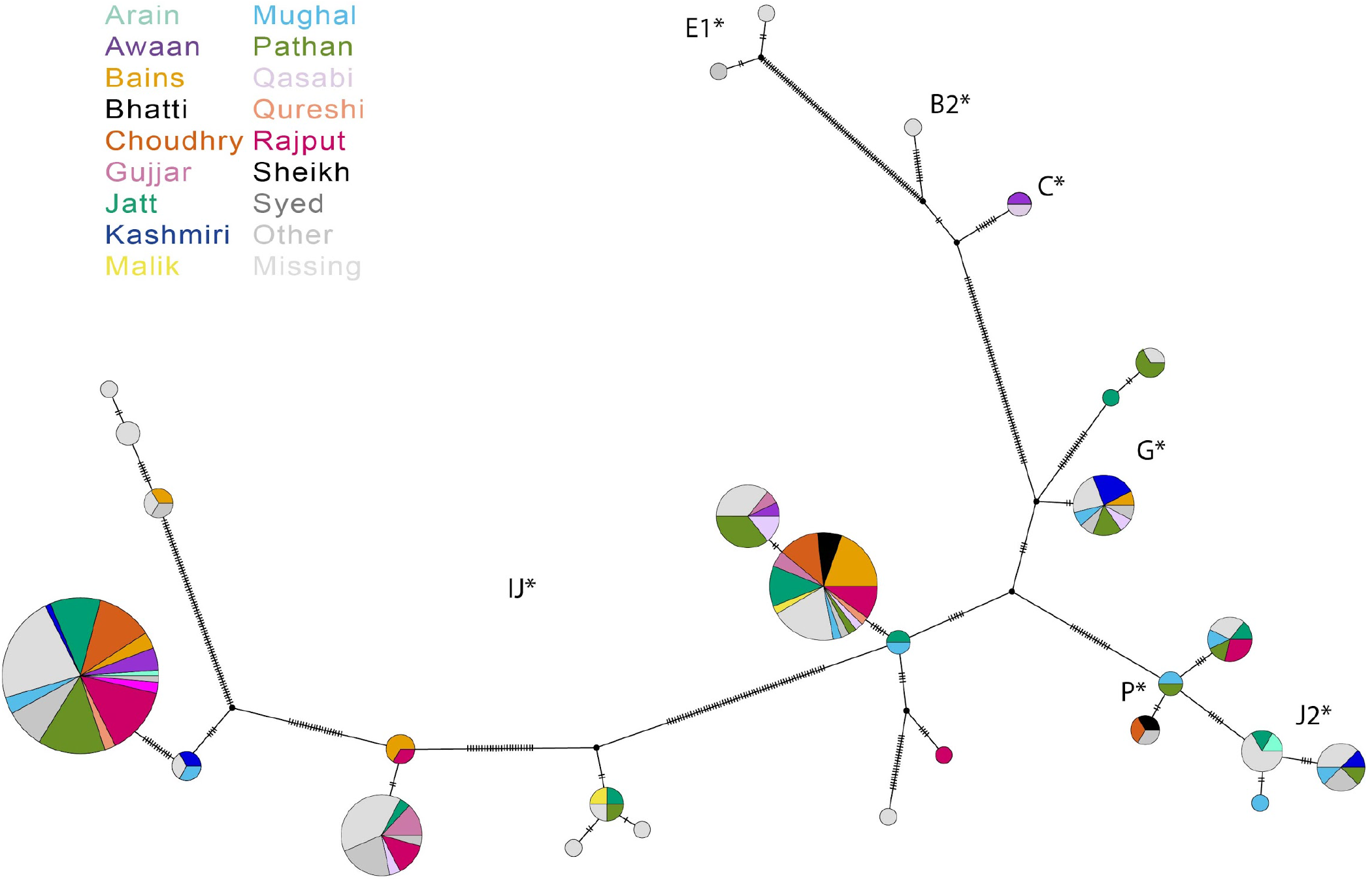
Y chromosome haplotype network based on median joining for 228 BiB Pakistani fathers, constructed from Y chromosome SNPs on the GSA chip. The area of the circles is proportional to the frequency of the haplogroup. Note that the branch length is *not* proportional to the genetic distance. The mutations separating each haplotype are indicated as hatch marks. The letters on the plot represent the different haplogroups.

**Supplementary Figure 11.**
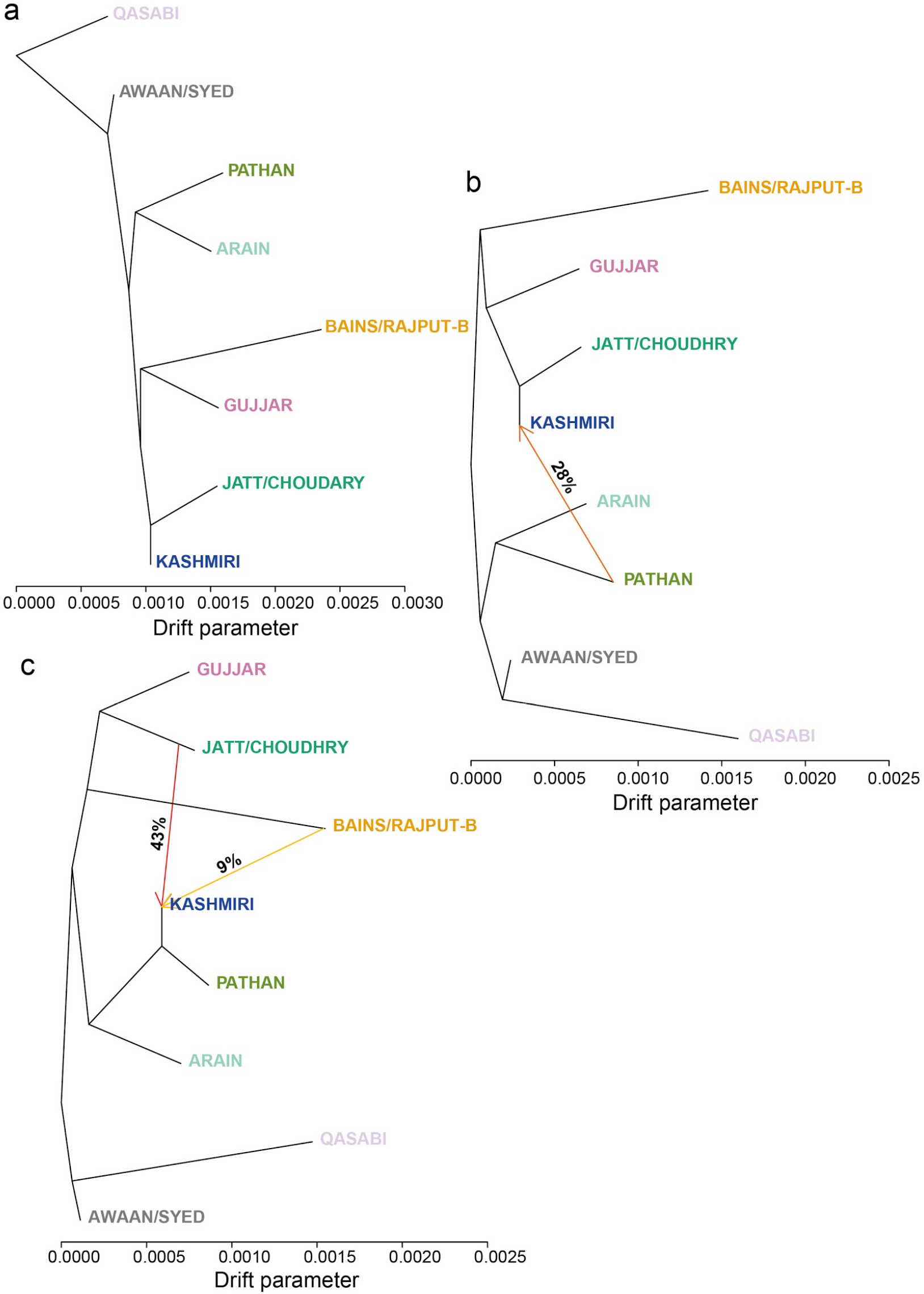
Inferred population trees with different mixture events based on Treemix analyses, using the homogeneous Pakistani subgroups defined with fineSTRUCTURE. The migration arrows are coloured and labelled according to their weight, which is correlated with the ancestry fraction shared. a) No migration edges allowed. b) One migration edge allowed. c) Two migration edges allowed. This could indicate, for example, that, according to the tree topology shown in (c), 43% of Kashmiri ancestry is derived from Jatt/Choudhry and 9% from Bains/RajputB. Note that the tree topology changes as migration edges are added, so one should not place too great a weight on it.

**Supplementary Figure 12.**
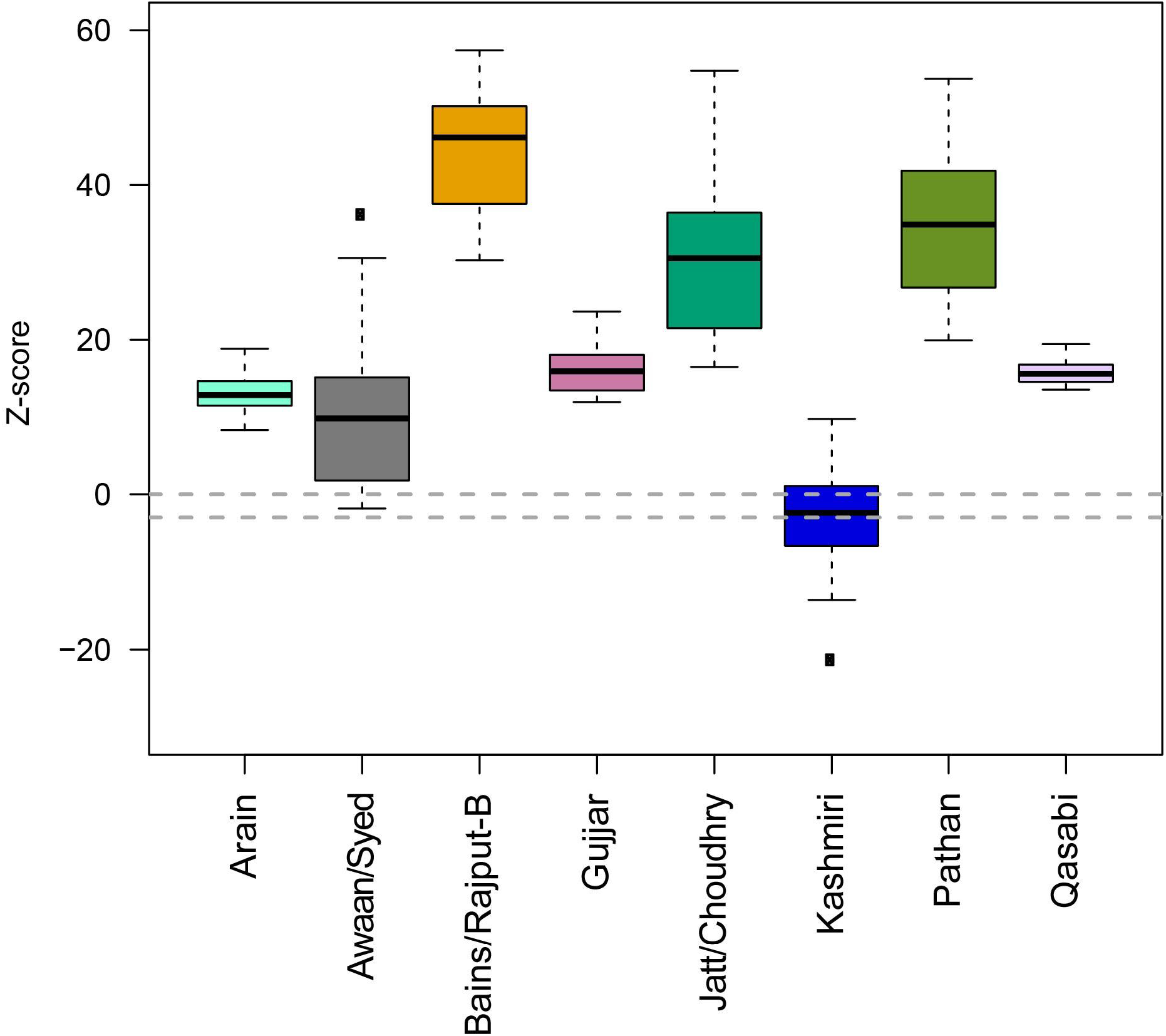
Boxplot of *f3*-statistics test of admixture, using all possible combinations of sources among the homogeneous Pakistani subgroups defined with fineSTRUCTURE. Dotted lines represent Z-score values of 0 and −3 which represent indication of putative admixture and significant admixture respectively. The only population that shows significant *f3* values is the Kashmiri.

**Supplementary Figure 13.**
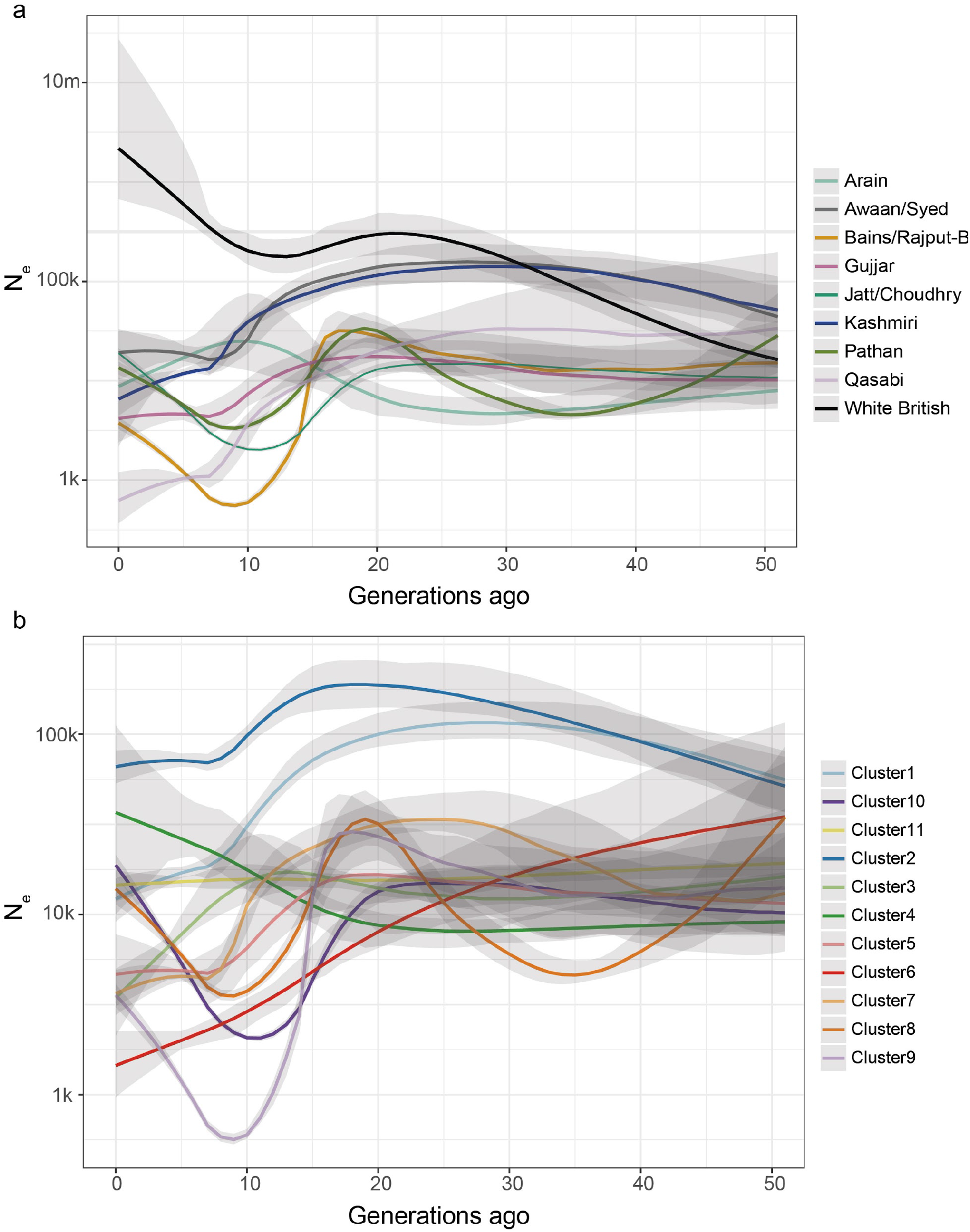
Effective population size (N_e_) changes through time estimated with IBDNe.The coloured lines indicate the mean estimate and the grey shading indicates 95% confidence intervals. a) Comparison of BiB White British and homogeneous Pakistani subgroups defined with fineSTRUCTURE. b) N_e_ estimates for all fineSTRUCTURE clusters (Figure 2).

**Supplementary Figure 14.**
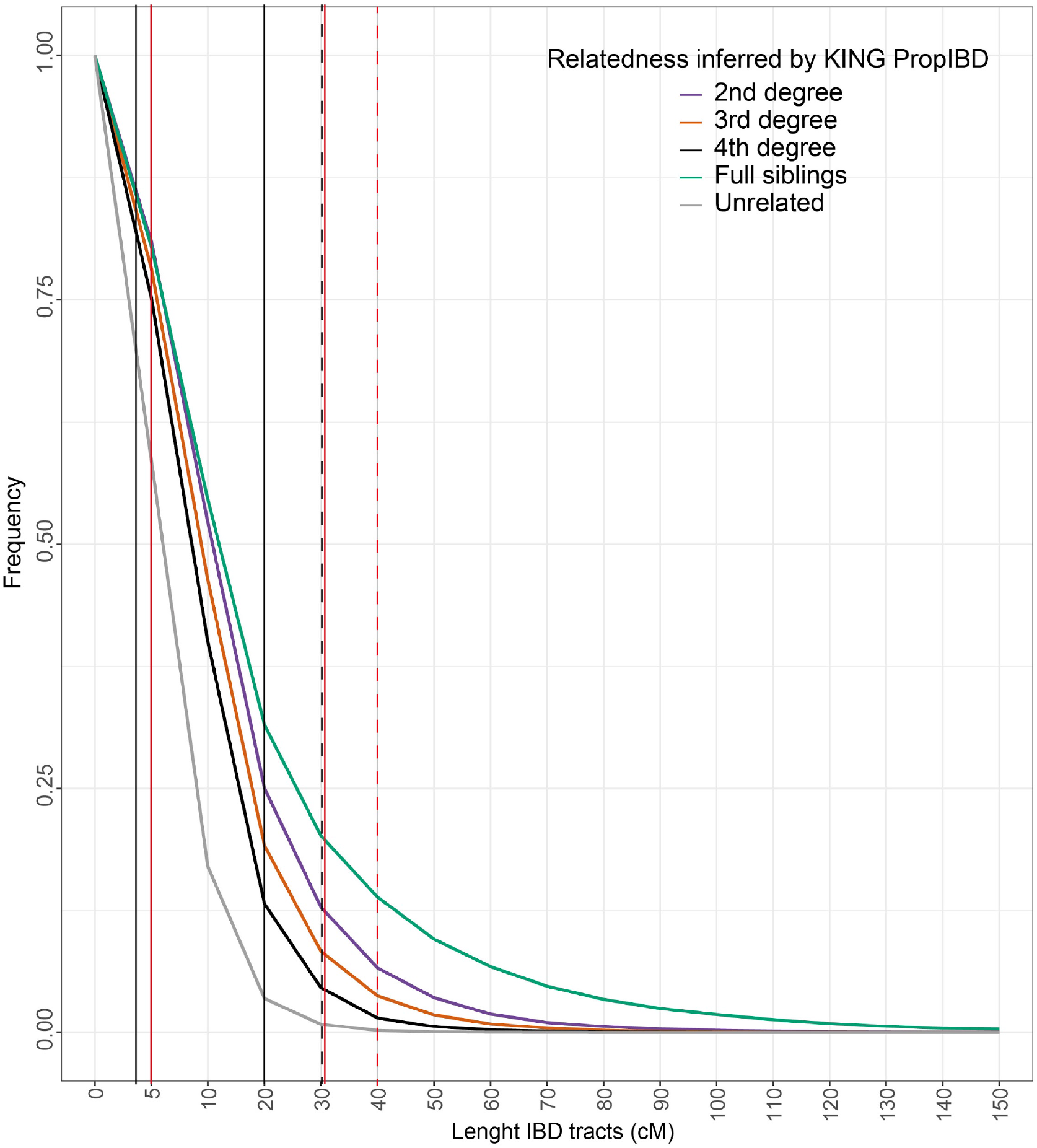
Cumulative distribution of IBD segments called by GERMLINE stratified by KING coefficient of relationship (PropIBD). The plot shows the proportion of individuals that share at least one segment greater than the length bin indicated on the x-axis, stratified by coefficient of relationship. The vertical lines represent the IBD thresholds used for IBD scores (see Methods, Figure 4): the dashed line represents the threshold used to define and exclude possible relatives and the full lines represent the range of IBD considered for the IBD score calculations. Red lines indicate those used for Figure 4, and black lines those used in Nakatsuka *et al*. and for Supplementary Figure 15a.

**Supplementary Figure 15.**
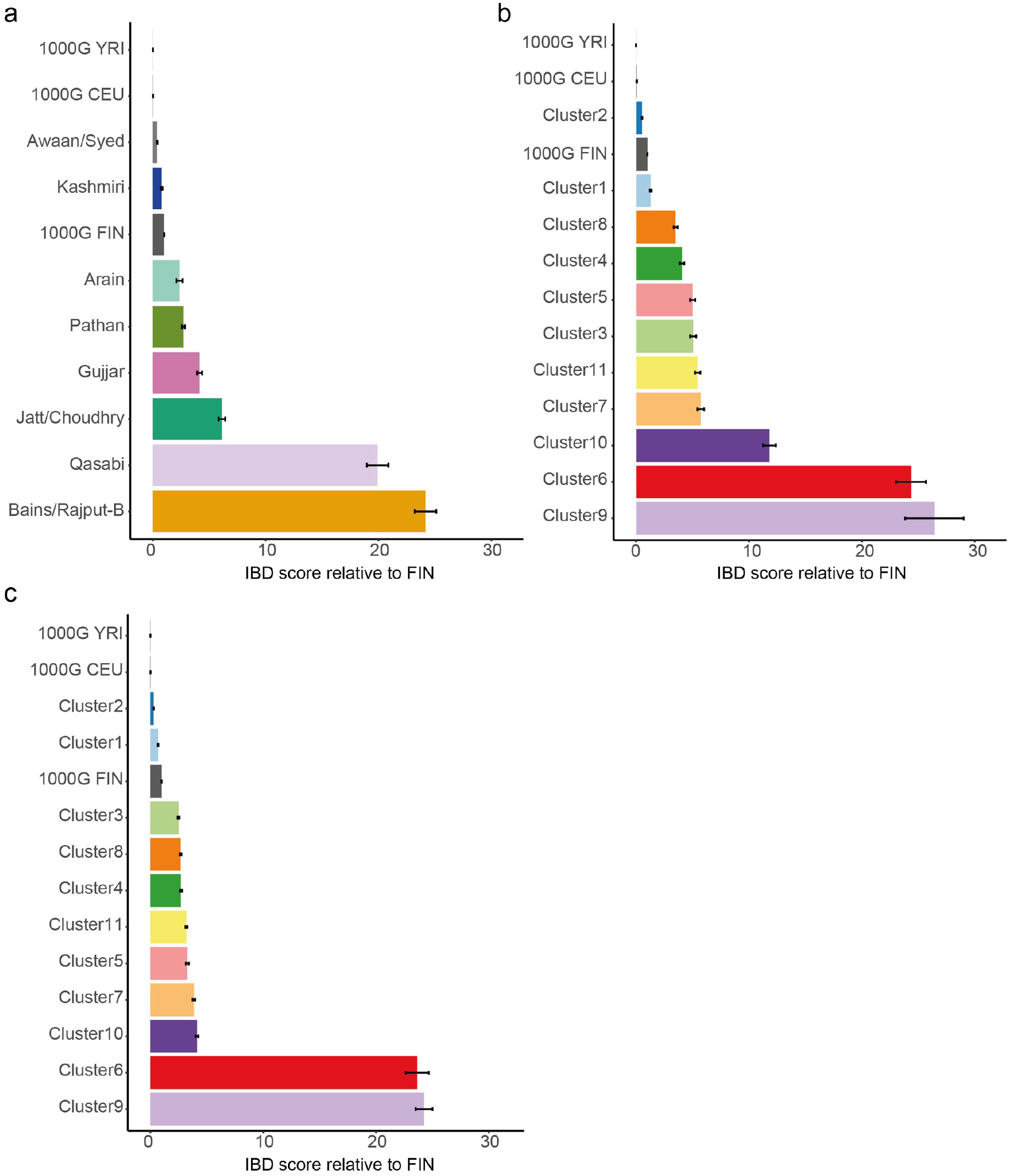
IBD scores calculated between individuals from the indicated group standardised by the value for the 1000 Genomes Finnish individuals. The error bars indicate the IBD score standard error. a) Homogeneous groups defined by fineSTRUCTURE considering the total length of IBD segments between 3 and 20cM, i.e. the same cutoffs used in Nakatsuka *et al*.^34^. b-c) All genetic clusters defined by fineSTRUCTURE, considering the total length of IBD segments between 5 and 30cM (i.e. the filtering used for Figure 4a) (b) or between 3 and 20cM (c).

**Supplementary Figure 16.**
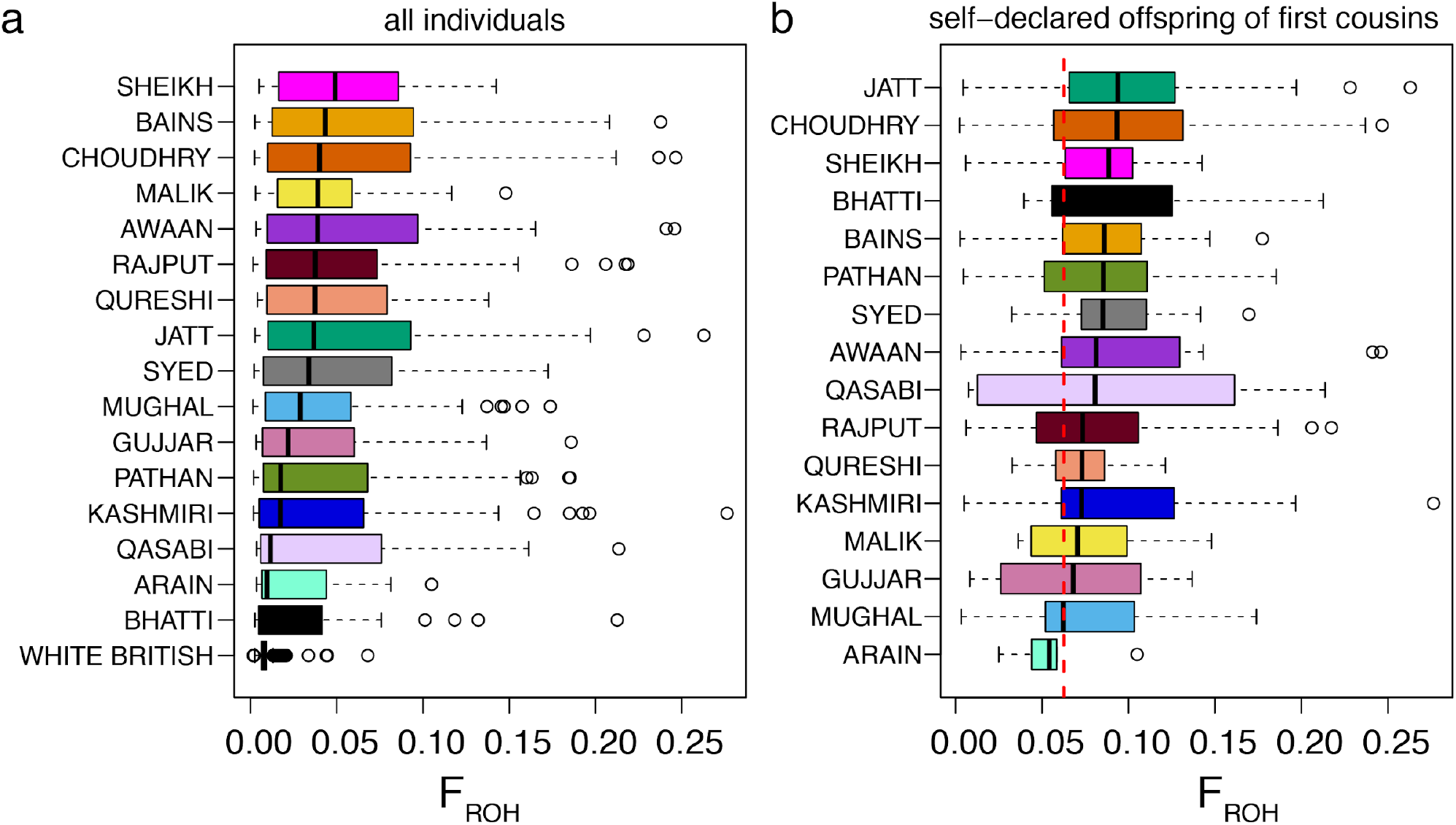
Boxplots showing the distribution of the fraction of the genome in regions of homozygosity (F_ROH_) in Pakistani versus White British mothers, with the Pakistanis broken down by self-reported subgroup. a) all individuals; b) individuals who selfreported as being offspring of first cousins. The red vertical line indicates the expectation for offspring of first-cousins whose parents are themselves unrelated.

**Supplementary Figure 17.**
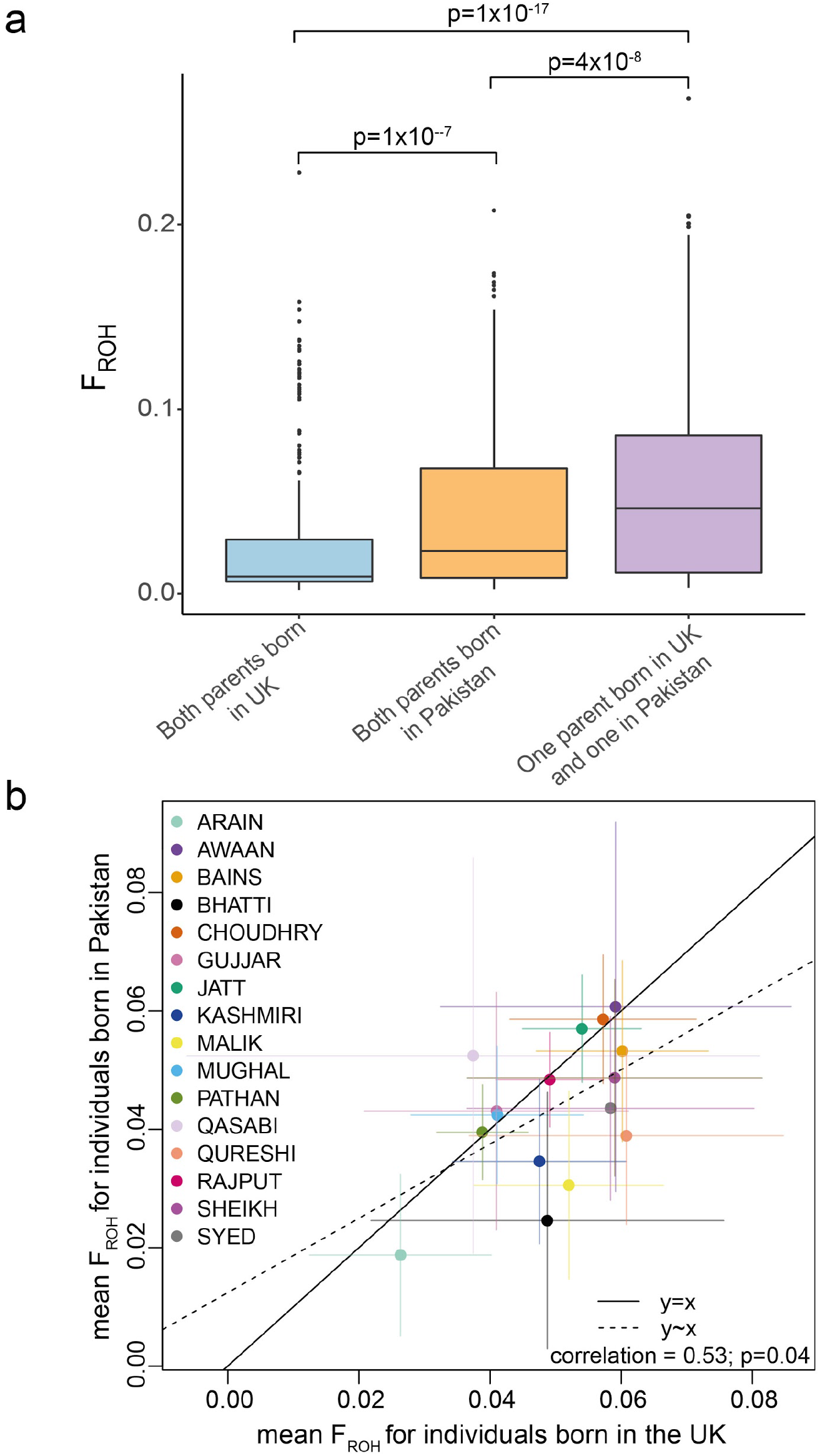
Patterns of F_ROH_ in BiB children and mothers according to parents’ or own birthplace. a) Boxplot of F_ROH_ for Pakistani children split by their parents’ birthplace. P-values are from Wilcoxon tests. b) Average F_ROH_ with 95% confidence intervals for Pakistani mothers born in Pakistan versus in the UK, split by self-declared subgroup. The slope for the line of best fit (dotted line) is 0.63 (standard error 0.27) (p=0.04). Only the Malik show a significant difference between those born in Pakistan versus the UK (p-value from one-sided t-test = 0.03).

**Supplementary Figure 18.**
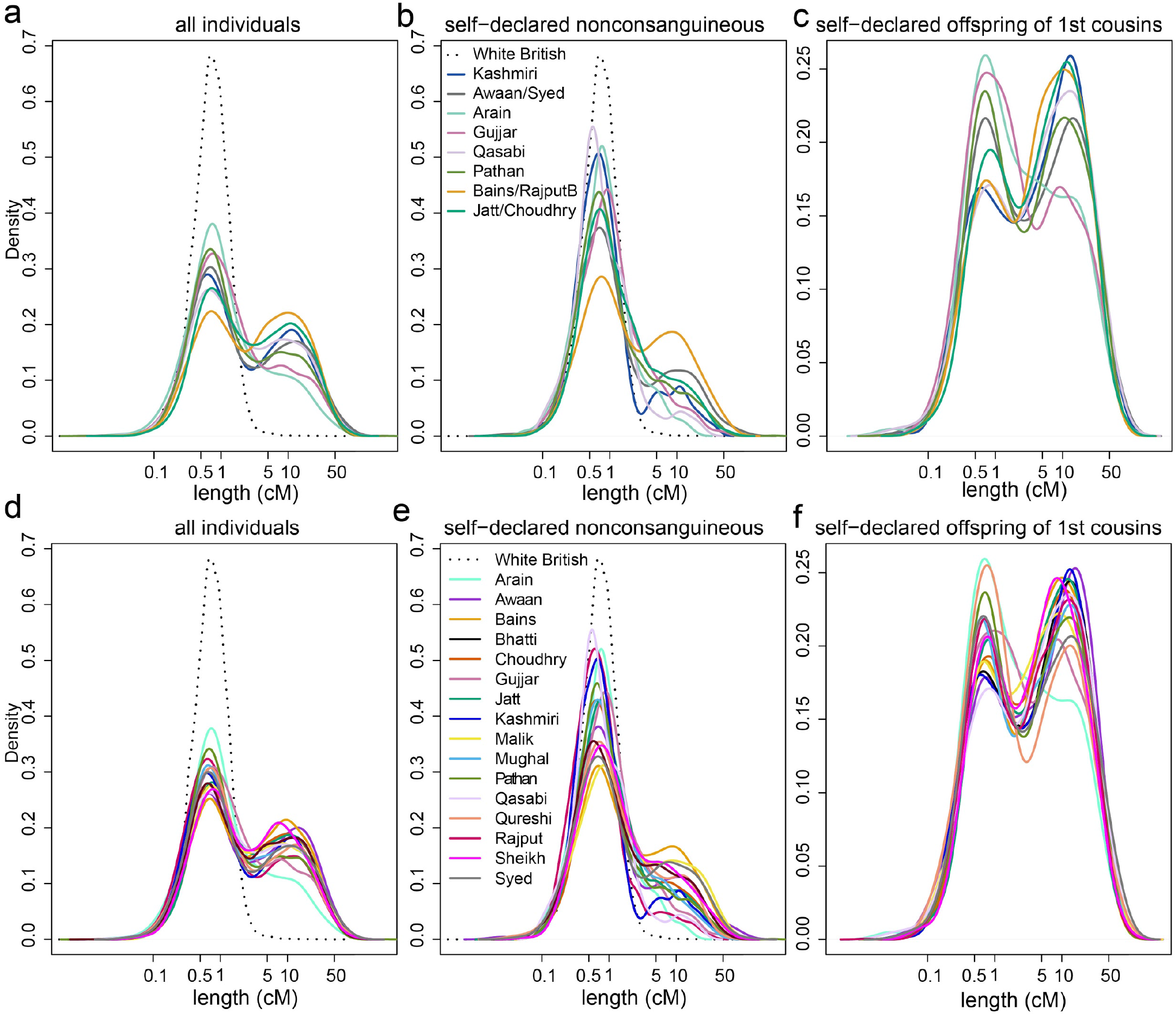
Distributions of the lengths of regions of homozygosity for White British mothers versus different subsets of the Pakistani mothers in BiB. a-c) Pakistani mothers split by homogeneous subgroup as defined using the fineSTRUCTURE clusters. d-f) Pakistani mothers split by self-reported groups. a,d) all mothers; b,e) all mothers self-declaring as having unrelated parents; c,f) all mothers who declared that their parents were first cousins.

**Supplementary Figure 19.**
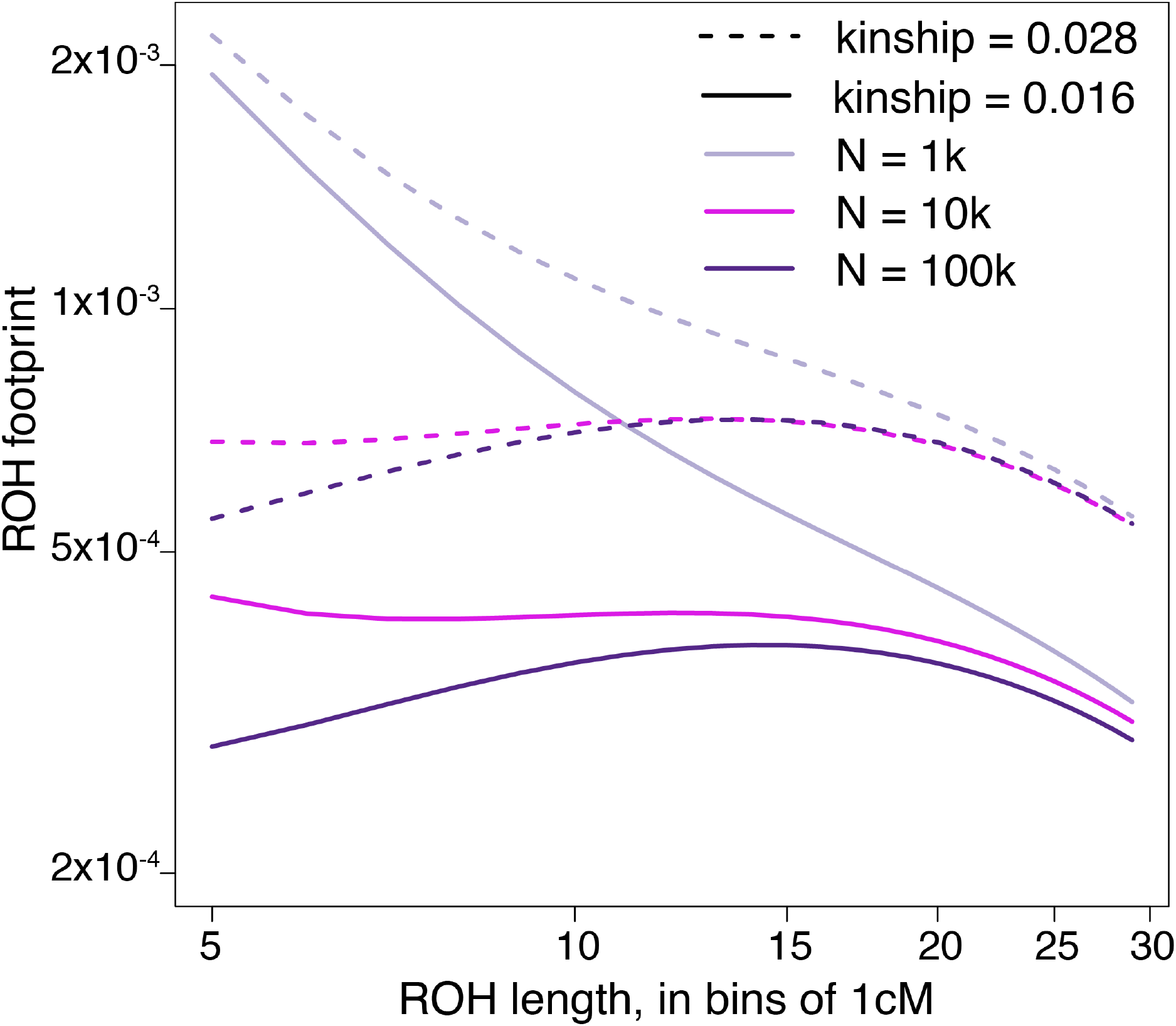
Impact of the effective population size and average kinship between spouses on the expected ROH footprint determined using the theory from ^62^ and Methods. The ROH footprint is the average fraction of the genome covered in ROHs of a given length. N = number of mating couples (i.e. N_e_/2). The average kinship values shown represent our naive estimates for Pathan (0.016) and Jatt/Choudhry (0.028) based on the fraction of the mothers who reported that their parents were first cousins, second cousins, or other relatives (which we assumed to be third cousins). The N_e_ has a major influence on the expected footprint of ROHs < 25cM.

**Supplementary Figure 20.**
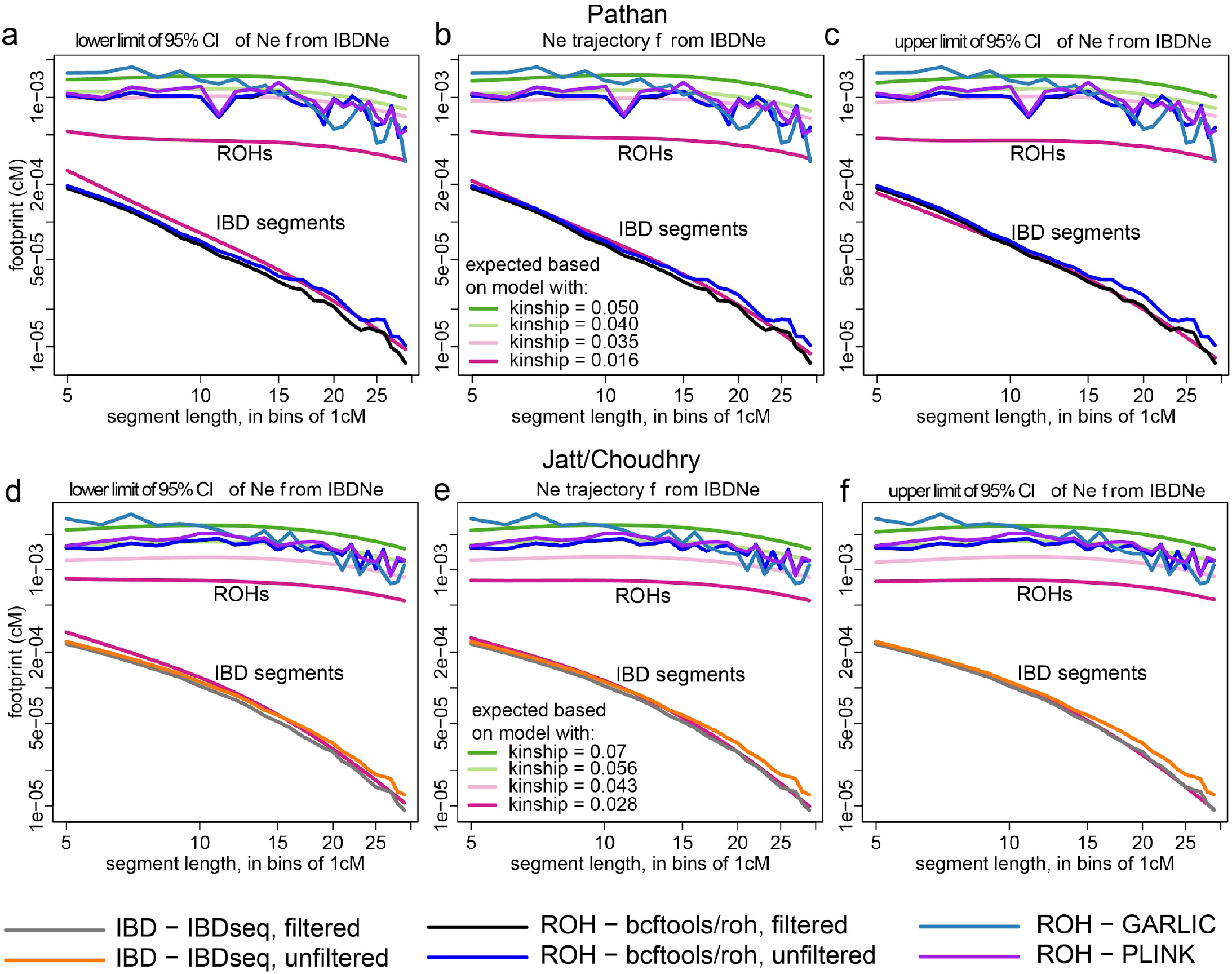
Observed ROH and IBD footprints compared to expectation under the model in ^62^. The footprint is the average fraction of the genome covered by segments of a given length interval. The top lines represent the ROH footprint and the bottom lines the IBD footprint. Points are plotted at the beginning of each 1cM interval. The expectation was determined using the indicated kinship values and either the point estimates of N_e_(t) from IBDNe (middle plots) or the lower or upper bound of their 95% confidence interval for *t* ≤ 50 (lefthand and righthand plots, respectively), then a constant N_e_ for *t* > 50. The IBDNe results used were for Pathan from fineSTRUCTURE Cluster 8 (a-c) or Jatt/Choudhry from Cluster 10 (d-f), and the corresponding observed footprints are plotted accordingly. We show the observed IBD footprint from IBDseq calls and the observed ROH footprint with bcftools/roh calls both with and without the filtering described in Methods, as well as the ROH footprint from PLINK and GARLIC ROH calls. Note that the expected IBD footprint does not depend on the kinship, so those lines are superimposed.

**Supplementary Figure 21.**
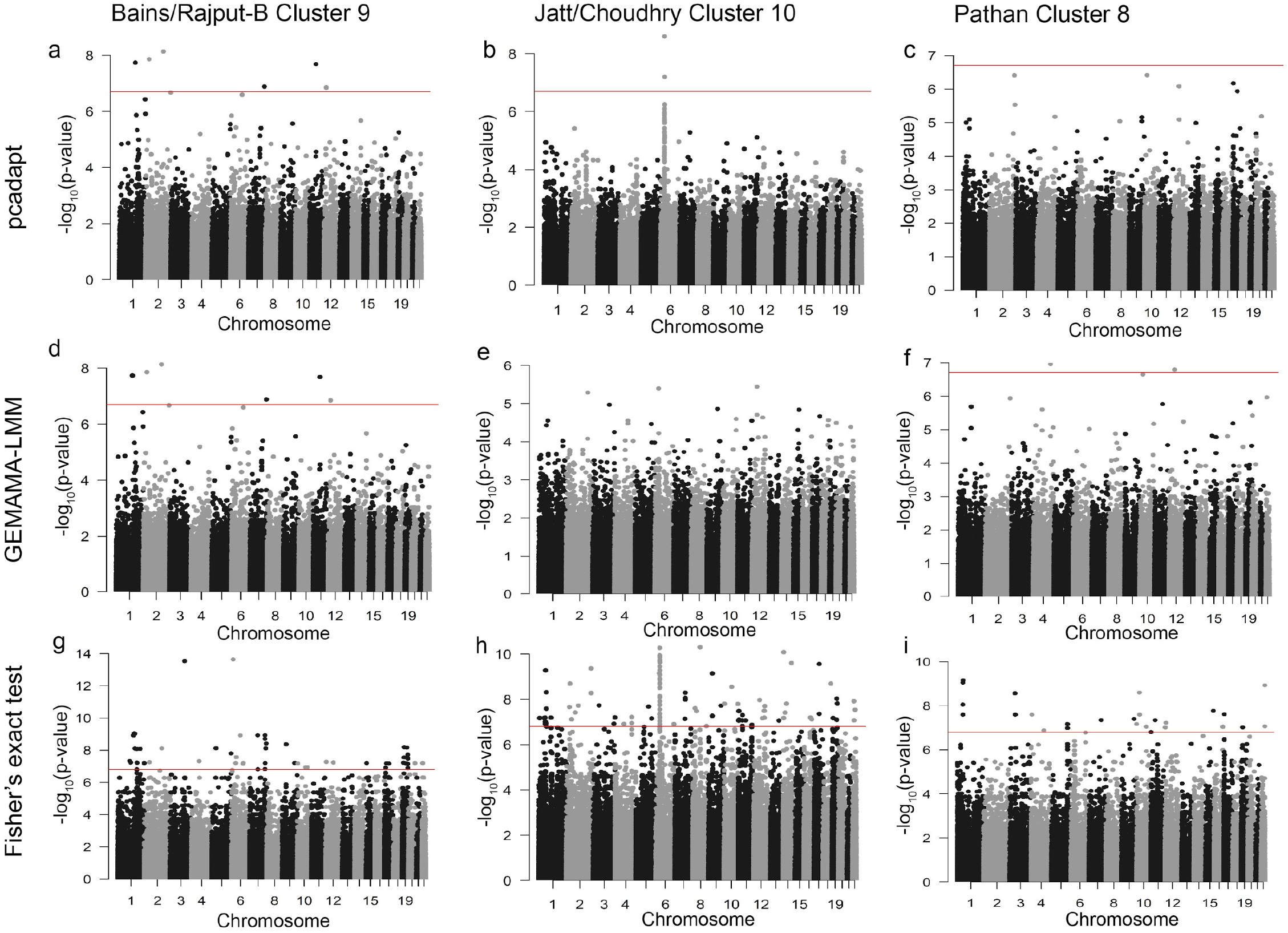
Analysis of variants with outlier frequencies in homogeneous subgroups. Manhattan plots of −log_10_(p-values) (y-axis) plotted against genomic coordinates (x-axis) from different methods for Bains/Rajput-B, Jatt/Choudhry and Pathan, as indicated in the column header. In each case, the index group was compared to individuals in Clusters 1-5 and Cluster 7 of the fineSTRUCTURE tree (Figure 2). The red line represents the Bonferroni corrected p-value (0.05/genome-wide number of variants tested). a,b,c) Results obtained applying the pcadapt method to the CoreExome data; d,e,f) Results obtained applying the GEMMA-LMM method to the CoreExome data; g,h,i) Results obtained applying the Fisher’s exact test to the whole-exome sequencing data

**Supplementary Figure 22.**
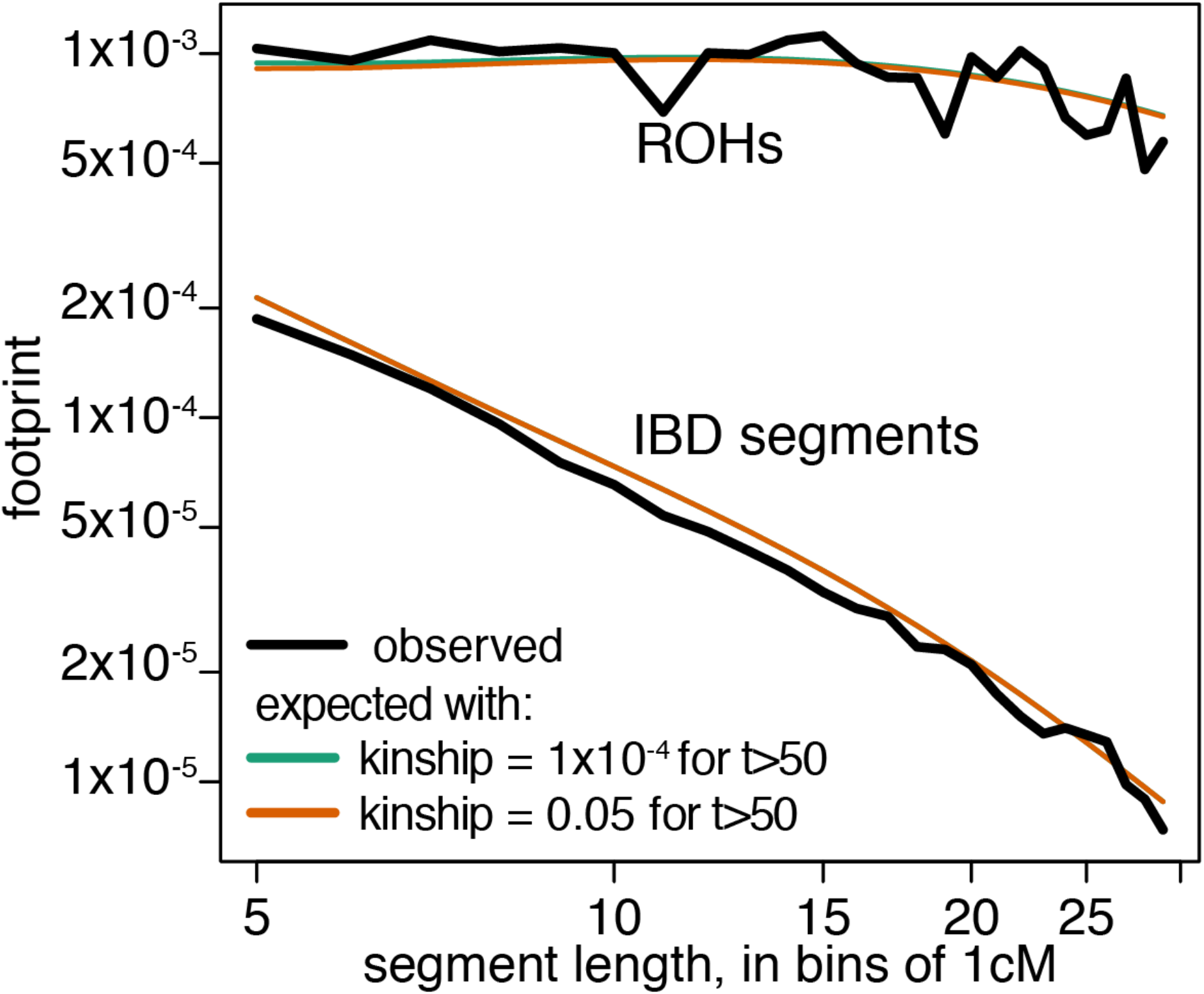
Effect of changes in historical consanguinity rates on the expected ROH and IBD footprints. The footprint is the average fraction of the genome covered by segments of a given length interval. The top lines represent the ROH footprint and the bottom lines the IBD footprint. Points are plotted at the beginning of each 1cM interval. The expectation was determined using the mean IBDNe estimate as N_e_(t) for Pathan from fineSTRUCTURE cluster 8 and an average parental kinship value, *k*, of 0.0354 for *t* ≤ 50, then using a constant N_e_ and the indicated *k* for *t* > 50. Note that the expected ROH footprint is barely altered when using *k*=1×10^−4^ versus *k*=0.05. The observed IBD footprint comes from IBDseq calls and the observed ROH footprint from bcftools/roh calls, with the filtering described in Methods.

**Supplementary Figure 23.**
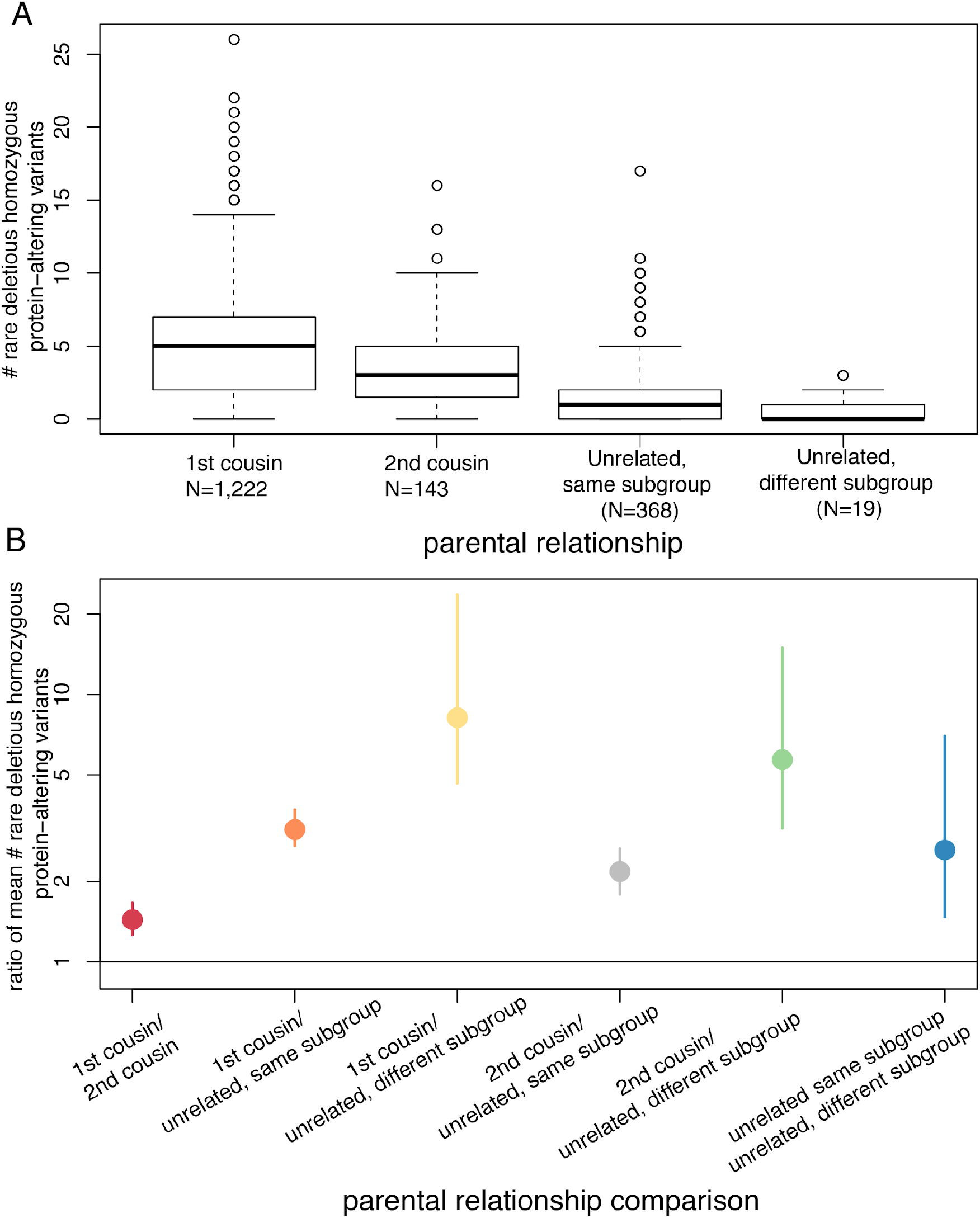
Effect of different levels of parental relationship on the number of rare deleterious homozygous protein-altering variants in the BiB mothers, inferred from exome-sequence data. A) Boxplot showing the distribution of the number of such variants per individual, stratified by self-reported parental relationship. b) Ratio of mean numbers of variants between groups of mothers who reported that their parents had the relationships reported on the X-axis. Error bars represent 95% confidence intervals from bootstrapping.

**Supplementary Figure 24.**
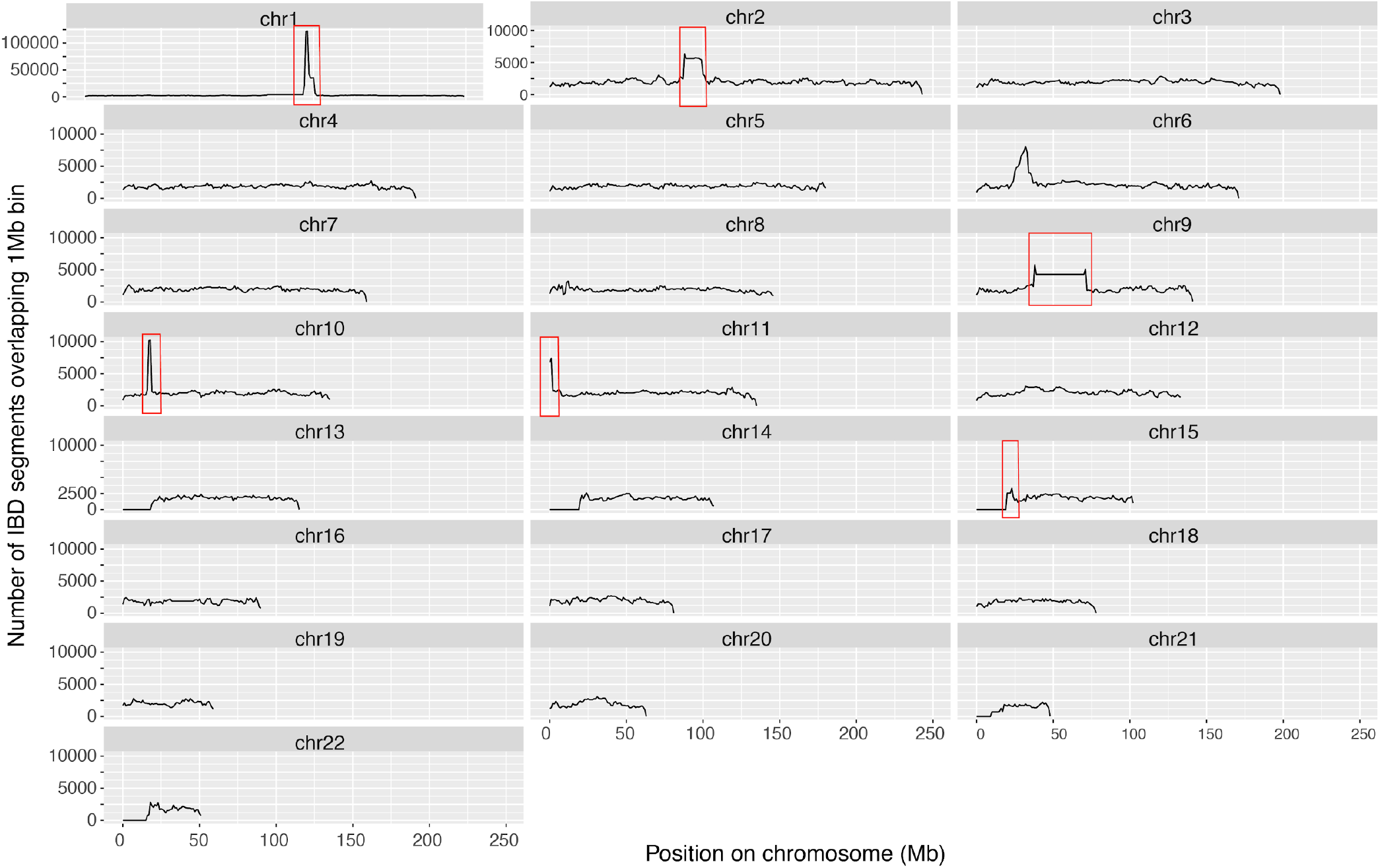
Likely artefacts in IBD segment calling with GERMLINE. The plots show the number of IBD segments called per 1Mb bin along each chromosome using the Bradford Pakistani mothers genotyped on the CoreExome chip. Note that the scale for chr1 differs from the one for the other chromosomes. The regions indicated with the red boxes appeared to be problematic in that they had a large number of identical IBD segments called over gaps. Several of these regions were also problematic in the IBDseq calling. We applied the filters described in the Methods to clean these data, and the results are shown in Supplementary Figure 25.

**Supplementary Figure 25:**
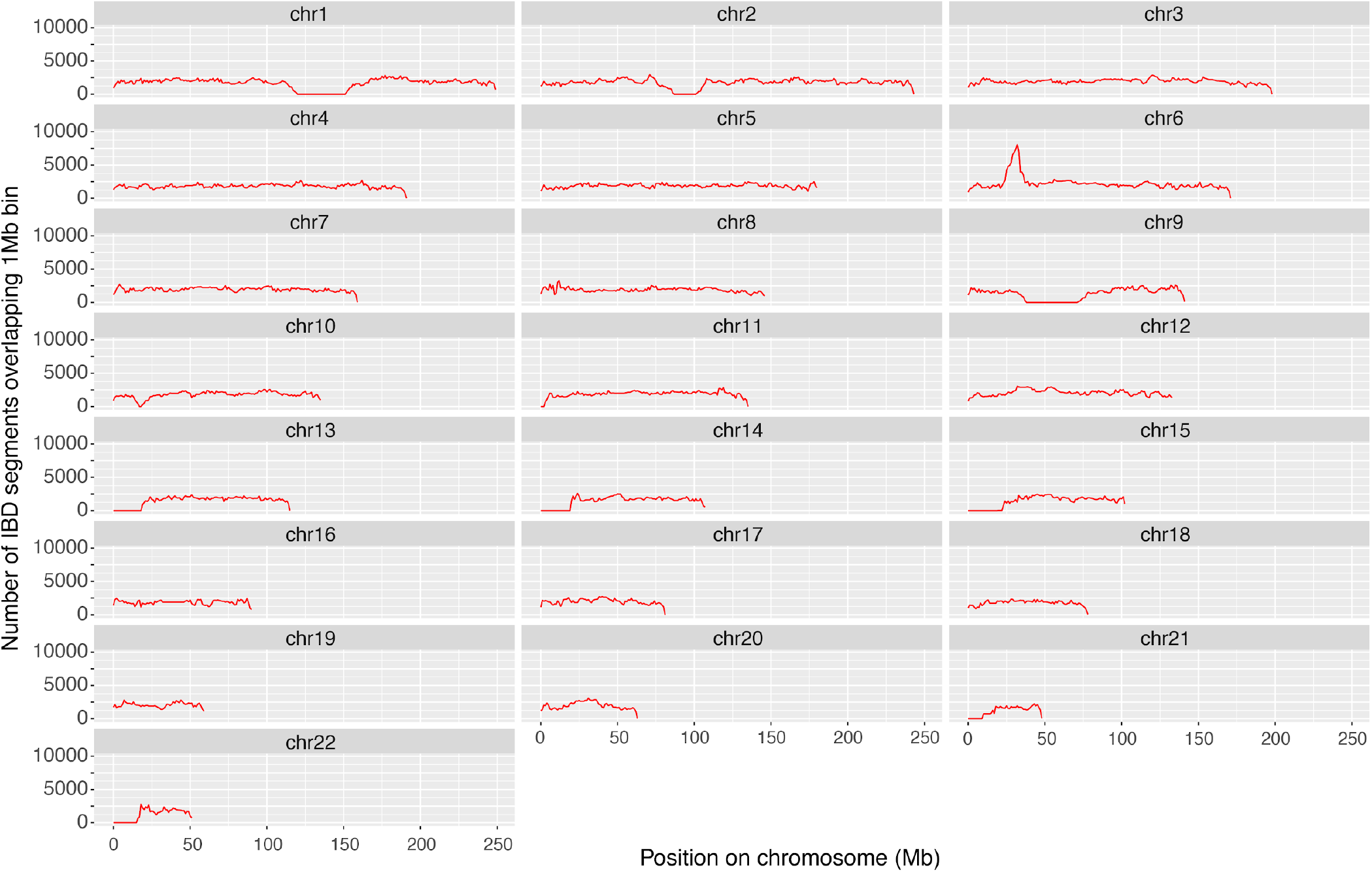
IBD segments called with GERMLINE, after filtering problematic regions as described in the Methods. The plots show the number of IBD segments remaining after filtering per 1Mb bin along each chromosome, using the Bradford Pakistani mothers genotyped on the CoreExome chip. Note that the peak on chr6 is around the HLA region, where there is likely to be increased IBD sharing due to selection. IBD segments overlapping the HLA region and centromeres were removed for the IBD score and IBDNe analyses.

**Supplementary Figure 26.**
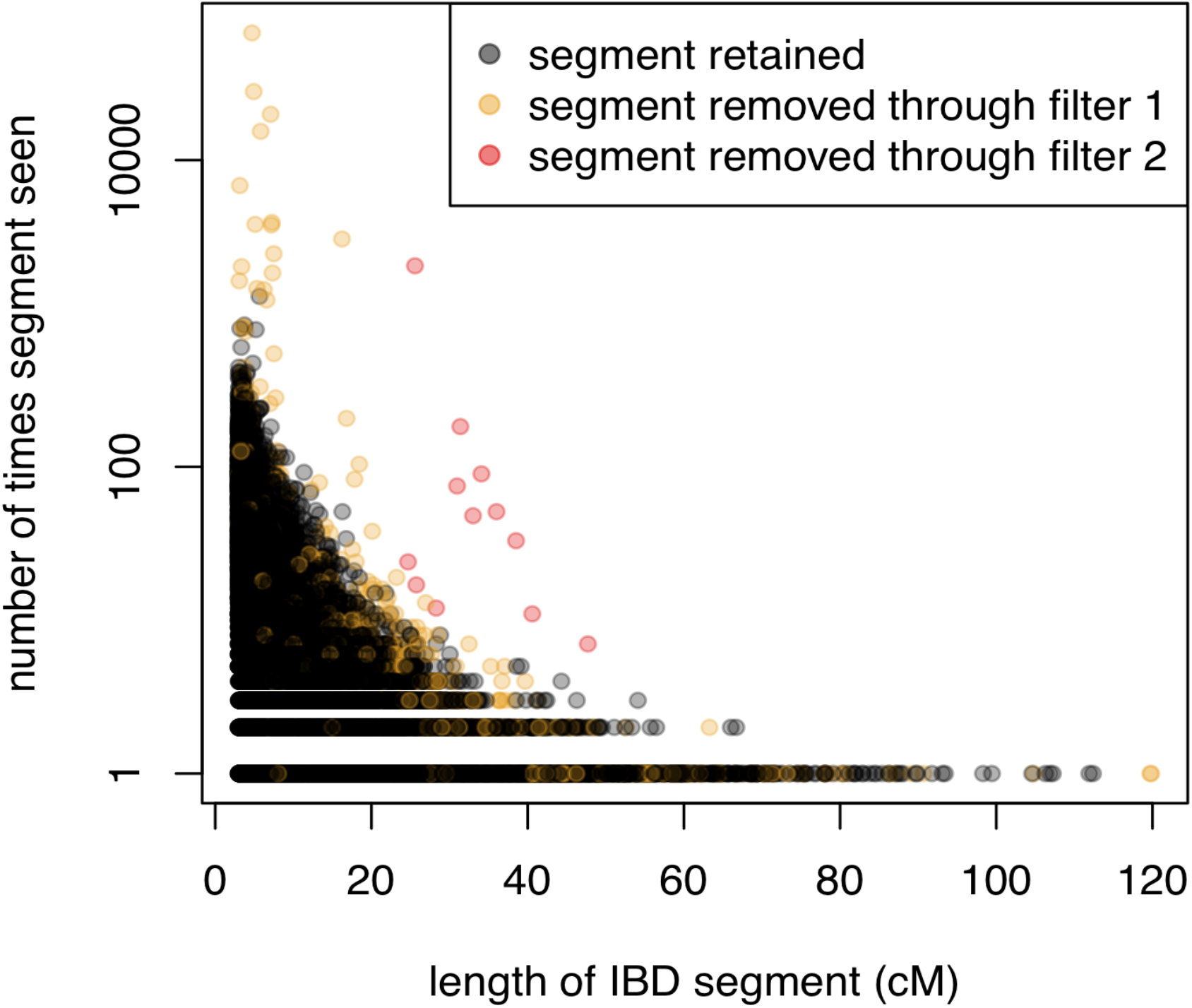
Filtering likely spurious IBD segments from GERMLINE (see Methods). The yellow points indicate segments in problematic regions that were apparent in the whole-genome plot shown in Supplementary Figure 24. The red points indicated segments that were removed in a further filter because they appeared to be called an unusually high number times given their length. These tended to overlap long gaps between SNPs and seemed likely to be artefacts.

**Supplementary Figure 27.**
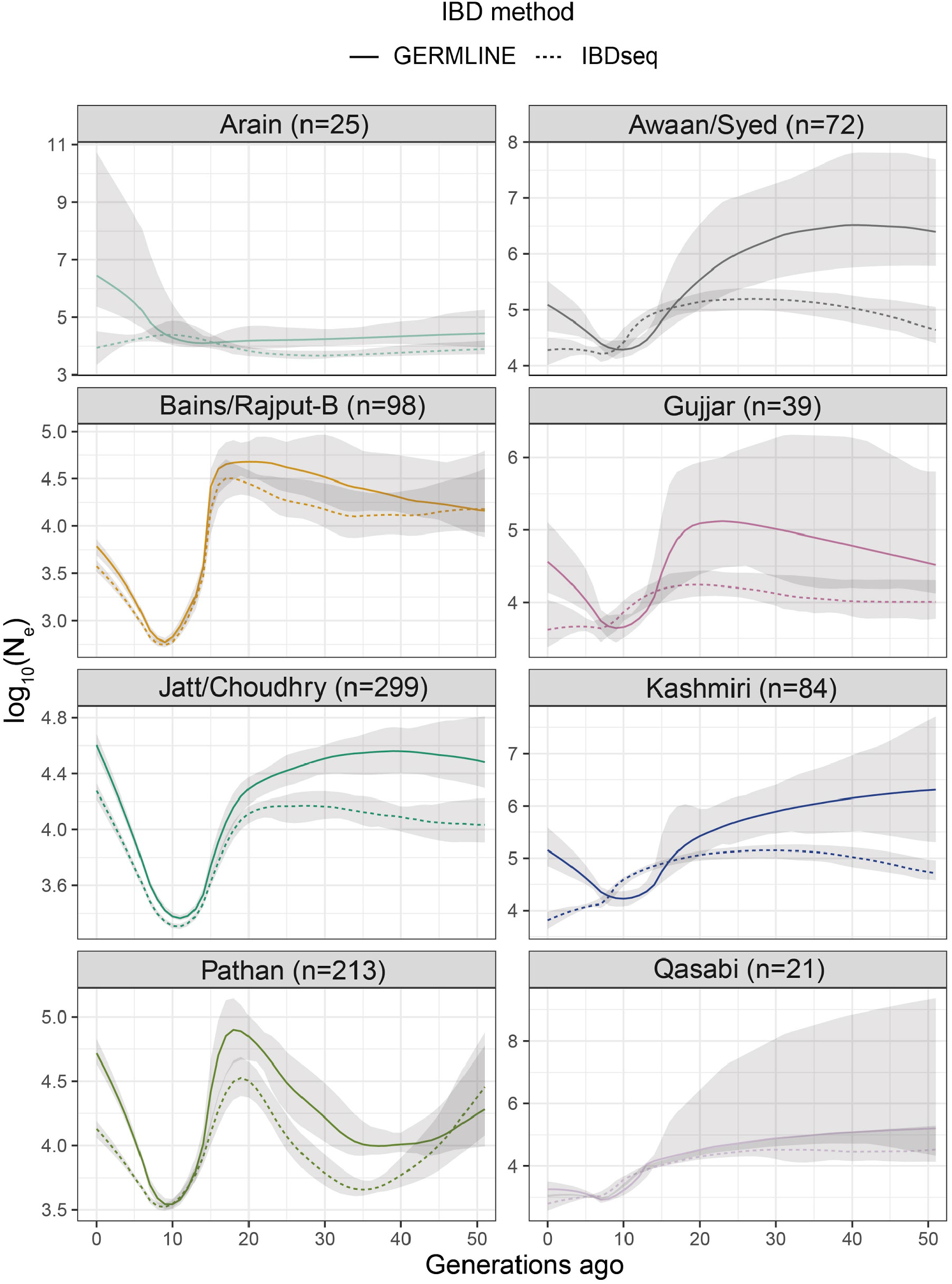
Change in effective population size (N_e_) through time estimated with IBDNe. The solid lines indicate the mean estimate using GERMLINE IBD output and the dashed line using IBDSeq output. The grey shading indicates 95% confidence intervals. The sample size for each subgroup is indicated in brackets.

## Supplementary Tables

**<u>Supplementary Table 1</u>**: Number of BiB samples by dataset and self-declared ethnicity.

**<u>Supplementary Table 2</u>**: Number of Pakistani and White British samples by dataset with participant information (mother/father/child), before outlier removal.

**<u>Supplementary Table 3</u>**: Number of Pakistani mothers and fathers (after sample QC) with self-declared subgroup information, before and after removing relatives using the KING PropIBD filter.

**<u>Supplementary Table 4</u>**: *f4*-statistics results for Bradford sub-groups reporting Arabic ancestry using other subgroups and Middle East populations. The *f4*-statistics were computed with the phylogeny *f4*(*W*, *X*; *Y*, Chimpanzee), where *W* represents the *biraderi* groups reporting Arabic ancestry (Qureshi, Sheikh or Syed), *X* are all the other Bradford subgroups and *Y* represents Middle East populations^39^. Positive values of *f4* statistics indicate gene flow between *W* and *Y*, but none of these are significant (Z<3 for all tests).

**<u>Supplementary Table 5</u>**: Y-chromosome haplogroups found in Pakistani fathers split by self-declared subgroup. The haplotype distribution listed in column B is from ^42,43^.

**<u>Supplementary Table 6</u>**: mtDNA haplogroups found in Pakistani mothers split by self-declared subgroup. The haplotype distribution listed in column B is from ^47,48,112^.

**<u>Supplementary Table 7</u>**: Pairwise *F*_ST_ values between each pair of homogeneous subgroups defined by fineSTRUCTURE, with their sample sizes.

**<u>Supplementary Table 8</u>**: Pairwise divergence times (in generations) between homogeneous subgroups defined by fineSTRUCTURE, inferred using NeON. Sample sizes are given in the second column. The numbers in brackets indicate 95% confidence intervals.

**<u>Supplementary Table 9</u>**: Pairwise divergence times (in generations) for all clusters defined by fineSTRUCTURE, inferred using NeON. The numbers in brackets indicate 95% confidence intervals.

**<u>Supplementary Table 10</u>**: Pairwise divergence times (in generations) within Cluster 9 (Bains/Rajput-B) and Cluster 10 (Jatt/Choudhry) defined by fineSTRUCTURE, inferred using NeON. The numbers in brackets indicate 95% confidence intervals.

**<u>Supplementary Table 11</u>**: Significant *f3*-statistics for Kashmiri using different pairs of the other homogeneous populations as source populations. The *f3* statistics were calculated in the form of *f3*(target, source 1, source 2). Significant tests (Z< −3) provide evidence that the target population is derived from an admixture of populations related to source 1 and source 2.

**<u>Supplementary Table 12</u>**: Rates of consanguinity as declared by BiB Pakistani mothers, split by self-declared subgroup. Note that the totals may not match those in Supplementary Tables 2 and 3 due to missing data in the questionnaire responses.

**<u>Supplementary Table 13</u>**: Rates of consanguinity as reported by BiB Pakistani mothers, split by homogeneous subgroup defined using fineSTRUCTURE. Note that the sample sizes may not match those given in Supplementary Table 8 due to missing data in the questionnaire responses.

**<u>Supplementary Table 14</u>**: Significant results from pcadapt and GEMMA-LMM, applied to CoreExome chip SNPs to identify variants at differing frequency in the largest homozygous subgroups compared to cluster 1-5+7. Only variants that passed multiple-testing correction in at least one of the two tests are shown. Fisher’s exact test p-values for the significant variants are reported. The allele frequencies given pertain to the alternative allele.

**<u>Supplementary Table 15</u>**: Results from the Fisher’s exact test applied to exome-sequence data to identify variants at differing frequency in the largest homozygous subgroups compared to cluster 1-5+7. Only variants that passed multiple-testing correction are shown. The allele frequencies given pertain to the alternative allele.

